# Metabolic Origins of Neurotoxic 1-deoxySphingolipids in Type 2 Diabetes

**DOI:** 10.1101/2025.04.25.650588

**Authors:** Adam Majcher, Par Bjorklund, Ales Holfeld, Aneta Cinakova, Alina Zurcher, Ermanno Malagola, Elkhan Yusifov, Sofia Kakava, Jerome Robert, Jennifer Härdfeldt, Katarina Hadova, Eva Kralova, Bodo Levkau, Alaa Othman, Giuseppe Lauria, Bengt Belgard, Bo Angelin, Thorsten Hornemann

## Abstract

Type 2 diabetes (T2D) and diabetic peripheral neuropathy (DPN) are associated with disruptions in sphingolipid (SL) metabolism, including an increased formation of neurotoxic 1-deoxysphingolipids (1-deoxySL). Here we report data from an untargeted proteomics, lipidomics and metabolomics profiling in plasma and skin samples of a carefully characterized T2D cohort and age-matched healthy controls. We investigated the association between plasma and skin amino acids and the sphingolipidome in blood and skin of T2D patients and several diabetic rodent models. We developed a hypothesis on how changes in the metabolism of the two amino acids relates to SL formation and DPN. To test this hypothesis and identify key enzymes responsible for the 1-deoxySL formation we developed stable isotope based SL flux assays, using UC13Glucose, 15NGlutamine, D4-Palmitic acid, D4-Alanine and D3N15-Serine as tracers. In combination with genetic interference approaches, we identified pathways that are responsible for shifting between the formation of 1-deoxySL and canonical SL in T2D. Furthermore, we verified these findings in vivo in several rodent models. This study links disturbances in amino acids, lipids, and protein homeostasis to DPN in T2D.

## Introduction

Over the past two decades, the global prevalence of adults with diabetes has more than tripled, reaching an estimated 537 million by 2021, with type 2 diabetes (T2D) accounting for over 90% of cases^1^. Diabetic polyneuropathy (DPN), the most common diabetes-related complication with a lifetime prevalence among T2D patients of approximately 50%, is now the most prevalent form of peripheral neuropathy overall^2^. DPN progressively impairs peripheral nerve function, manifesting in a range of clinical symptoms including neuropathic pain, loss of sensation and weakness. DPN is also a major cause of skin ulcerations and limb amputations and significantly impacts patients’ quality of life^3^. The pathophysiological mechanisms underlying DPN remain poorly understood, and to date, no effective targeted therapy is available. Hyperglycemia-related mechanisms are suspected to exert neurotoxic effects on neurons and glia cell through increased formation of reactive oxygen species and pro-inflammatory advanced glycation end products. Despite aggressive treatment of hyperglycemia, the incidence of DPN remains high^4^. Current management approaches focus on lifestyle modifications, pain management, and intensive glucose control. Peripheral nervous system is rich in lipids but their role in the development of peripheral neuropathy has been neglected for a long time.

Sphingolipids (SL) – a highly diverse class of bioactive lipids with signaling functions and important roles in forming functional cellular membranes have been previously implicated in peripheral neuropathies^5^. SL biosynthesis starts with the formation of a sphingoid base, the fundamental structural element of all SLs. Sphingoid base formation is the first rate-limiting step in the SL *de novo* synthesis and is catalyzed by the enzyme Serine Palmitoyl Transferase (SPT). Typically, SPT catalyzes the conjugation of L-serine (L-ser) with palmitoyl-CoA to produce (keto)sphinganine, which is then metabolized through several intermediate steps to ceramides and complex SLs^6^. Besides L-Serine, SPT can also utilize other amino acid substrates, primarily L-alanine (L-ala), forming an atypical SL class known as 1-deoxysphingolipids (1-deoxySL)^7^. 1-deoxySL lack the essential C1-OH group of canonical SL, which prevents their conversion into complex SL, but also their terminal degradation via the canonical SL catabolic pathway^8^. 1-deoxySL exert time- and dose-dependent neurotoxicity, both in vitro and in vivo ^9–15^. In the monogenic disease Hereditary Sensory and Autonomic Neuropathy Type 1 (HSAN1) mutations in SPT lead to a permanent shift in substrate preference from L-ser to L-ala which results in pathologically increased 1-deoxySL levels, and causes progressive axonopathy^16–20^. Also native SPT forms significant amounts of 1-deoxySL when availability of L-ser is restricted^12,21,22^. Recent studies have shown that limiting L-ser results in peripheral sensory deficits in insulin resistant mice while L-ser supplementation has been shown to effectively lower 1-deoxySL levels in HSAN1 and preserve nerve function in both mice and humans ^23–25^.

Clinically, DPN and HSAN1 share many characteristics and are both associated with increased 1-deoxySL formation, as demonstrated in animal and human studies. In T2D, 1-deoxySL plasma levels correlate inversely with intraepidermal nerve fiber density (IENFD), which is a measure of axonal damage^26–28^. In a diabetic rat model, L-ser supplementation lowered 1-deoxySL and improved sensory nerve function without affecting hyperglycemia^25^. Recently, Handzlik et al. demonstrated in a murine model that L-ser restriction, combined with obesity, insulin resistance, and dyslipidemia, leads to increased 1-deoxySL formation and sensory neuropathy^23^. Conversely, L-ser supplementation as well as inhibiting SPT activity, alleviated the neurotoxic effects^23^. However, differences in the L-ala and L-ser metabolism in T2D and their relation to 1-deoxySL formation and DPN remain unclear.

## Results

### Multiomics analysis of human T2D skin revealed accumulation of 1-deoxySL

Diabetic peripheral neuropathy (DPN) primarily affects skin sensation, where it typically starts as a small fiber neuropathy^2^. To better understand the molecular mechanisms underlying this early onset condition, we performed an untargeted multiomics analysis of human T2D skin biopsies, including lipidomics, metabolomics and proteomics LC-MS/MS based analysis, to identify metabolic alterations contributing to intradermal small fiber loss.

We analyzed the plasma and skin lipidome between clinically precisely characterized T2D patients (N=38) and sex- and age-matched healthy controls (N=39) (Fig. 1a). The Lipidomics profile of skin biopsies revealed 379 annotated lipid species, of which 68 were sphingolipids. While most other lipid classes remained unchanged between groups, 1-deoxySL were identified as the most significantly increased lipids in T2D skin (Fig. 1 b,c, Extended Fig. 1a,e). This suggests that 1-deoxySL accumulation is a metabolic feature of T2D skin. Given the known neurotoxic properties of 1-deoxySL, the increase might contribute to the loss of intraepidermal small fiber and the progression into incident DPN. In matching plasma samples, we identified 349 altered lipid species, including 72 sphingolipids. Consistent with prior reports, we observed a significant increase in plasma 1-deoxySL in T2D (Extended Fig. 1b,c), while other abundant SL such as sphingomyelins (SM) and hexosyl ceramide (HexCer) were decreased. Ceramide and sphingosine-1-phosphate levels remained unchanged (Extended Fig. 1d).

**Figure 1.**
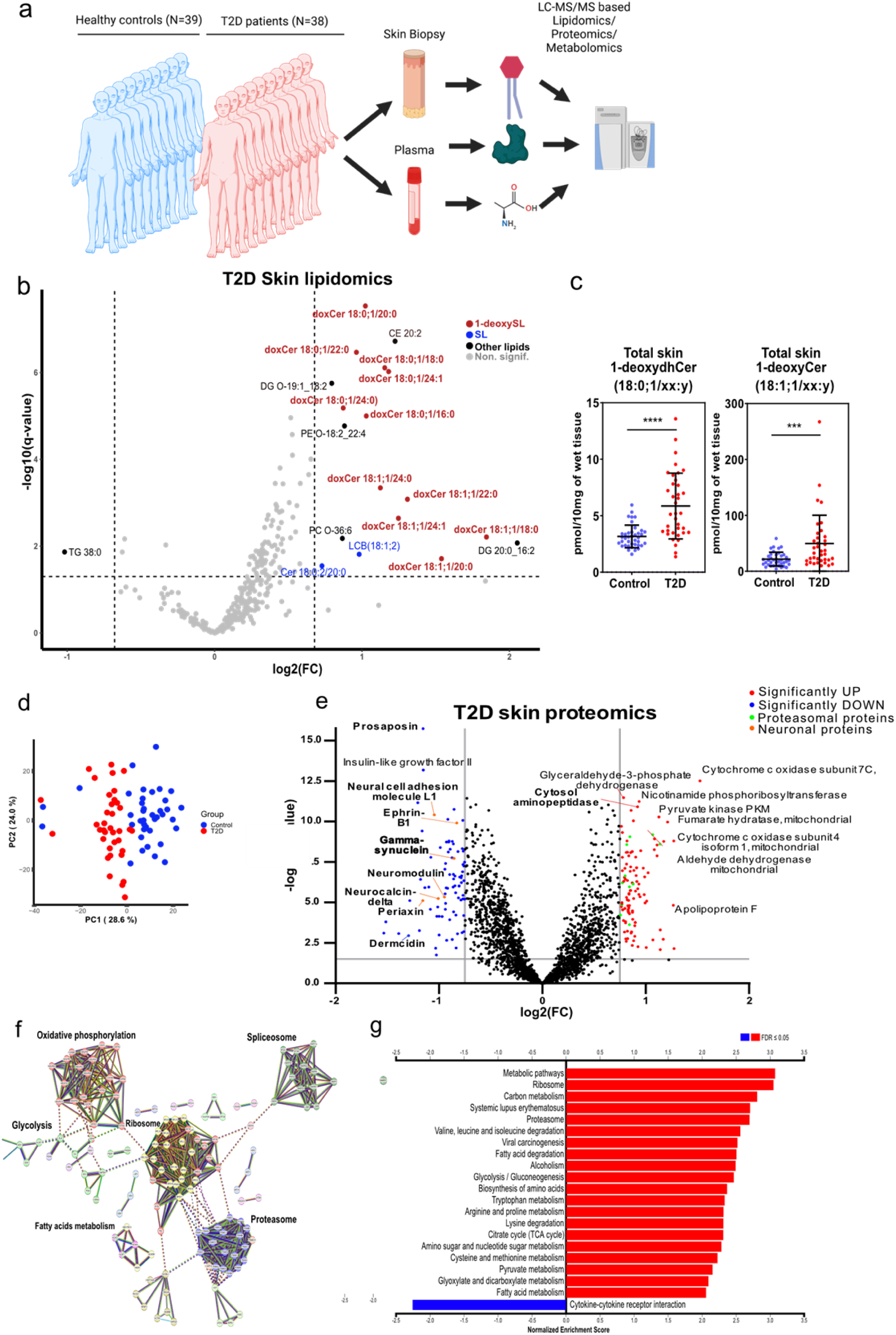
Multiomics analysis of human T2D skin. **a** Schmatic representation of sampling and analysis of our Cohort of 38 T2D patients and 39 age and sex-matched healthy controls. Plasma and skin punch biopsies were analysed with an LC-MS/MS based untargeted lipidomics and proteomics as well as in a targeted metabolomics approach. **b** Volcano plot showing lipidomics data of T2D diabetic skin compared to healthy controls (cutoff |log2FC| ≥ 0.75, p-value ≤ 0.05). 1-deoxySL (red) and canonical SL (blue) are indicated. **c** Total skin 1-deoxyDHCer (m18:0) and 1-deoxyCer (m18:1) per 10mg of wet tissue. Data are represented individually and as mean ± SD. P-values indicate the significance of the group comparisons (two-way Student’s unpaired t-test); *=p<0.05, **=p<0.01, ***=p<0.001, ****=p<0.0001. **d** Principal component analysis (PCA) of the skin proteome (2,309 quantified proteins) shows a distinct clustering between T2D and control groups, indicating disease-specific proteomic signatures. **e** Volcano plot showing proteomics data of T2D skin (N=38) compared to healthy controls (N=39). Cutoffs: log2FC ≥ 0.75, q-value ≤ 0.05. Proteins enriched (red) and reduced (blue) in T2D skin are indicated. Proteasomal proteins are represented in green, neuronal proteins in orange. **f** STRING analysis of the upregulated proteins in the skin of T2D. The highest confidence interval of interaction (0.9) was applied. Subsequently, MCL clustering (inflation parameter=3) was used. Clusters were colored using the KEGG database. **g** Gene set enrichment analysis (GSEA) based on log2FC values from differential analysis of the skin proteomics between T2D patients (N=38) and healthy controls (N=39). Pathways were parsed by the KEGG database.

To further explore metabolic alterations in T2D skin, we also performed untargeted proteomics, identifying 2,309 proteins. Unsupervised principal component analysis (PCA) revealed a distinct clustering of the skin proteome between the T2D and control group (Fig. 1d). Differential expression analysis identified 111 significantly up- and 84 down regulated proteins in T2D skin (cutoff |log2FC| ≥ 0.75, q-value ≤ 0.05) (Fig. 1e). A STRING pathway analysis showed five major clusters of upregulated proteins, including fatty acid metabolism, glycolysis, oxidative phosphorylation, spliceosome, ribosome and proteasome pathways (Fig. 1f). Notably, the upregulation of ribosomal and proteasomal factors suggests increased protein turnover in T2D skin. Conversely, several neuronal markers, including prosaposin, γ-synuclein, and ephrinB1, were decreased, suggesting neuronal loss in T2D skin (Fig. 1e). Gene set enrichment analysis (GSEA) revealed a significant upregulation of pathways related to ribosome and proteasome activity, carbon metabolism, amino acid metabolism, central energy metabolism (pyruvate metabolism, citrate cycle, glycolysis/gluconeogenesis), and fatty acid degradation (Fig. 1g). These findings suggest that metabolic reprogramming in T2D skin primarily affects energy metabolism and protein homeostasis, which may contribute to the pathogenesis of DPN. Altogether, these results demonstrate 1-deoxySL accumulation and proteome-wide metabolic dysregulation in T2D skin, supporting their potential role in the progression of diabetic neuropathy.

**Supplementary Figure 1.**
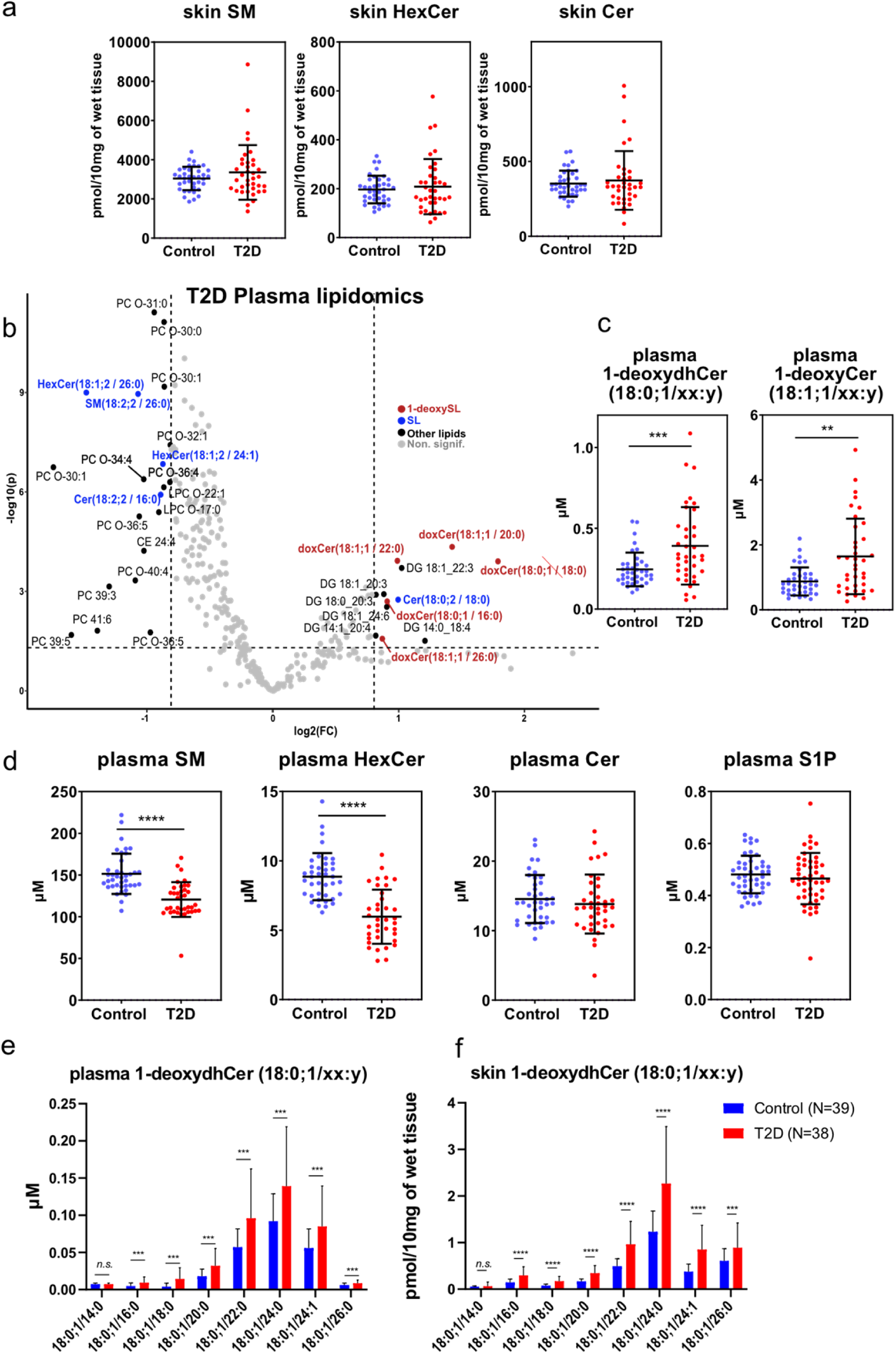
Multiomics analysis of human T2D skin. **a** Total levels of main Sphingolipid subclasses in skin of 39 Healthy and 38 Diabetic skin biopsies measured by untargeted LC-MS/MS analysis. **b** Volcano plot showing lipidomics data of T2D diabetic plasma compared to healthy controls (cutoff |log2FC| ≥ 0.75, p-value ≤ 0.05). **c** Total plasma 1-deoxyDHCer (m18:0) and 1-deoxyCer (m18:1) levels. **d** Total levels of main Sphingolipid subclasses in corresponding plasma of 39 Healthy controls and 38 T2D patients measured by untargeted LC-MS/MS analysis. Data are represented as individual values and mean ± SD. P-values indicate the significance of difference between the study groups (two-way Student’s unpaired t-test). **e** Individual 1-deoxydhCer (m18:0/xx:y) with different acyl chain lengths in skin and **f** plasma. Bar plots data are represented as mean ± SD, P-values indicate the significance of difference between the study groups (two-way, Student’s unpaired t-test with Welch correction), multiple testing was corrected using a two-stage step-up method (Benjamini, Krieger and Yekutieli). *=p<0.05, **=p<0.01, ***=p<0.001, ****=p<0.0001.

### Alanine-to-Serine ratio dictates SPT activity and 1-deoxySL formation

Next, we sought to address the underlying mechanisms of 1-deoxySL formation in T2D. Mutations in SPT associated with HSAN1 cause a permanent substrate shift from L-ser to L-ala, leading to excessive 1-deoxySL formation (Figure 2a). To compare the impact of L-ala and L-ser on 1-deoxySL formation, we expressed SPTLC1^WT^ and the most common HSAN1 causing variant SPTLC1^C133W^ in SPTLC1 deficient HEK 293 cells. The ala/ser ratio was modulated by adding increasing concentrations of D4-Alanine at constant concentrations of isotope labelled D3N15Serine (0.5mM). The incorporation of the labeled amino acid substrates into the de-novo formed SL and 1-deoxySL was analyzed by LC-MS/MS. As reported, HSAN1 mutants have an increased activity with D4-Alanine compared to WT SPT. However, also the native SPT formed significant amounts 1-deoxySLs+D3 as a result of increasing D4-Alanine availability (Figure 2b).

**Figure 2.**
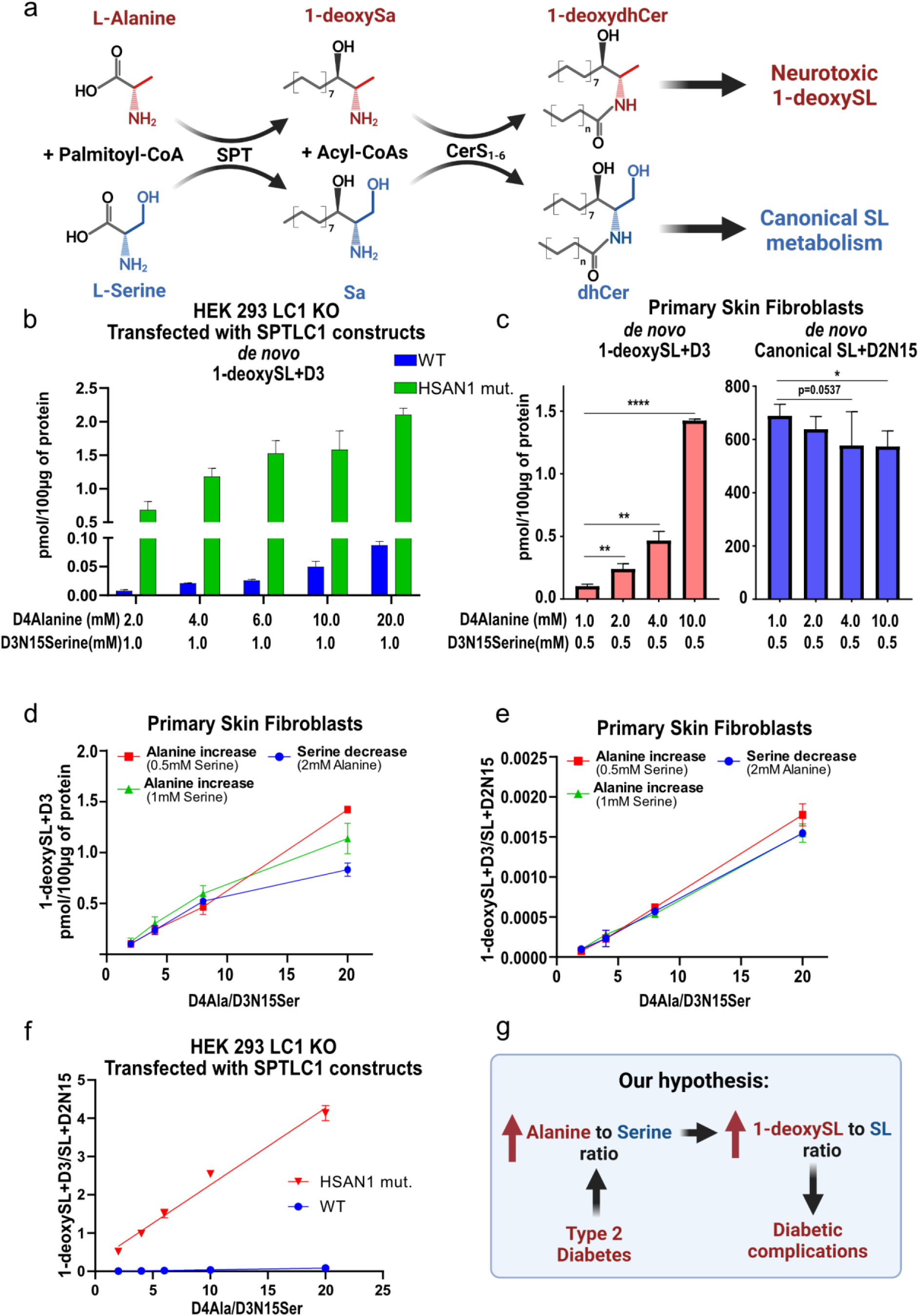
Alanine-to-Serine ratio dictates SPT activity and 1-deoxySL formation. **a** De novo synthesis of 1-deoxySL and canonical SL. Typically Serine-Palmitoyl transferase (SPT) utilizes L-Serine for the synthesis of canonical SL. The use of L-Alanine by the SPT forms neurotoxic 1-deoxySL which are associated with sensory neuropathies. 1-deoxySphinganine (1-deoxySa, doxLCB 18:0;1), Sphinganine (Sa, LCB 18:0;2), 1-deoxydihydroCeramides (1-deoxydhCer, doxCer 18:0;1/xx:y), dihydroCeramides (dhCer, Cer 18:0;2/xx:y). **b** Total levels of de novo synthesized 1-deoxySL+D3 after increasing d4-L-Alanine (D4Alanine) with stable concentration of D3-15N-L-Serine (D3N15Serine) in HEK 293 cells overexpressing SPTLC1^WT^ or the most common HSAN1 associated mutation SPTLC1^C133W^. **c** Total levels of de novo synthesized 1-deoxySL+D3 and canonical SL+D2N15 after increasing D4-L-Alanine (D4Alanine) with stable concentration of D3-15N-L-Serine (D3N15Serine) in primary skin fibroblasts. **d** Total de novo formed 1-deoxySL+D3 amounts and **e** 1-deoxySL+D3 to SL+D2N15 ratio (1-deoxySL+D3/SL+D3) in primary skin fibroblasts as a response to increasing D4Alanine to D3N15Serine ratio (D4Ala/D3N15Ser). Note that ratios can be increased by either raising Alanine or decreasing Serine. Red and green represent variable L-ala against constant L-ser concentrations (0.5mM and 1mM D3N15Ser, respectively). Blue represents decreasing L-ser levels at constant L-ala concentration (2mM D4Alanine). **f** 1-deoxySL+D3/SL+D3 ratio in HEK293 either overexpressing SPTLC1^WT^ or the HSAN1 mutation SPTLC1^C133W^ at increasing D4Ala/D3N15Ser. Levels of de novo formed 1-deoxySL and SL were analyzed using LC-MS/MS as indicated in the methods. All data are represented as mean ± SD. P-values indicate the significance of difference between the groups (two-way Student’s unpaired t-test); *=p<0.05, **=p<0.01, ***=p<0.001, ****=p<0.0001. **g** Our hypothesis postulating that increased L-Ala-to-L-Ser ratio (ala/ser) leads to increased 1-deoxySL-to-canonical SL ratio (1-deoxySL/SL) in T2D.

Analogous results could be observed when exposing primary human fibroblasts with increasing concentrations of D4-Alanine while maintaining a constant D3N15Serine concentrations, which revealed a strong positive correlation between L-alanine and 1-deoxySL formation, which was paralleled by a proportional reduction in canonical SL synthesis (Fig. 2c). Thus, these results suggest that alongside genetic alterations, the relative substrate availability is a critical modulator SPT activity.

We further expanded on these analyses to include other in vitro model system, such as liver (Huh7) and renal (HK2) cell models (Supplementary Fig. 2 a,d). These experiments confirmed that 1-deoxySL formation is an intrinsic property of the SPT enzyme and independent of cell and tissue type. Moreover they revealed that, rather than by the absolute amino acid concentrations, SPT mediated 1-deoxySL formation is dictated by the ratio between L-ala and L-ser. In line with this hypothesis, regardless of which substrate was modulated, positive increase of ala/ser ratio systematically associated with increased 1-deoxySL levels and reduced canonical SL synthesis (Fig. 2d, Supplementary Fig. 2c). Moreover, when directly correlating the ala/ser and 1-deoxySL/SL ratios, we observed a surprisingly wide range of linear relationship that was independent of absolute amino acid levels (Fig. 2d, Supplementary Fig. 2c,e). Analogous consideration could be drawn from the analysis of SPTLC1^C133W^ activity, albeit at an increased slope confirming that ala/ser ratio is the key factor dictating 1-deoxySL formation (Figure 2e).

Given these findings, we hypothesized that T2D patients exhibit an altered ala/ser ratio balance, leading to increased 1-deoxySL accumulation (Fig. 2g). To test this, we performed LC-MS/MS amino acid profiling in our T2D cohort, revealing significantly reduced L-ser and strongly elevated L-ala concentrations in T2D plasma, resulting in a markedly increased ala/ser ratio (Extended Fig. 2a,b). As predicted, ala/ser correlated positively with total 1-deoxySL levels and negatively with canonical SL levels (Extended Fig. 2c,d), leading to a strong association between ala/ser and 1-deoxySL/SL ratios in T2D patients (Extended Fig. 2c,d). This relationship was further validated by analysing plasma samples from an independent cohort of 685 individuals. In addition to the previous clinical data, these results confirmed a significant correlation between plasma ala/ser and 1-deoxySL/SL (Extended Fig. 2f), supporting the hypothesis that disruptions in the amino acid homeostasis could drive 1-deoxySL formation in T2D.

Importantly, the skin 1-deoxySL/SL ratio was also associated with clinical incident DPN. Comparing T2D patients with and without neuropathy (Extended Fig. 2g) revealed a significantly higher 1-deoxySL/SL ratio in the group with incident DPN (Extended Fig. 2h), reconfirming that DPN is associated with increased 1-deoxySL/SL ratio. Overall, these results establish ala/ser ratio as a critical determinant of 1-deoxySL synthesis and associated with DPN.

**Extended Figure 2.**
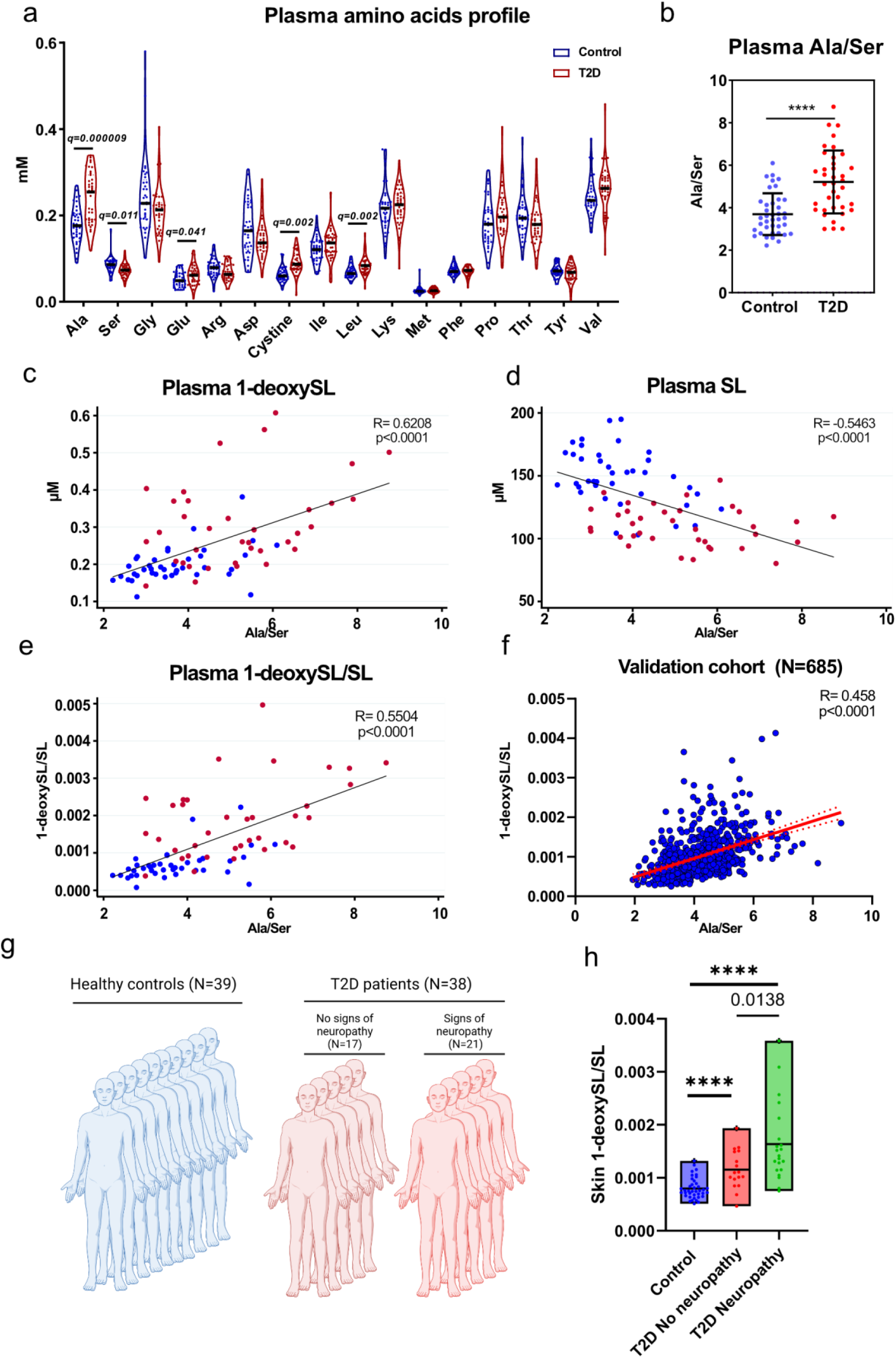
Alanine-to-Serine ratio dictates SPT activity and 1-deoxySL formation. **a** Violin plots showing plasma amino acid levels in T2D (N=38) and healthy controls (N=39). Glycine (Gly), Glutamate (Glu), Arginine (Arg), Aspartate (Asp), Isoleucine (Ile), Leucine (Leu), Lysine (Lys), Methionine (Met), Phenylalanine (Phe), Proline (Pro), Threonine (Thr), Tyrosine (Tyr), Valine (Val) were measured by targeted LC-MS/MS metabolomics as described in the methods.. Statistical significance was determined according to Mann-Whitney. Correction for multiple testing was done according to Benjamini, Krieger and Yekutieli **b** The ala/ser in plasma of T2D (N=38) compared to healthy controls (N=39). P-value indicates the significance between group comparisons (Student’s unpaired t-test), ****=p<0.0001 **c** Correlation of plasma ala/ser with total 1-deoxySL and **d** canonical SL **e** Correlation between plasma ala/ser and the 1-deoxySL/SL ratio in our cohort. Correlations were calculated according to Pearson (N=77, red T2D, blue Healthy). **f** Association between 1-deoxySL/SL and ala/ser in plasma samples of our T2D and an independent population based cohort (“CoLaus” N=685). Correlations were calculated using Pearson. **g** Cohort stratification according to the NDR neuropathy score. NDR was determined as described in methods. **h** Box plot showing the 1-deoxySL/SL in the T2D group according to their NDR status and in comparison with the healthy controls. **T2D No Neuropathy** reflects patients with T2D and NDR status=1 (N=17), and **T2D Neuropathy** with T2D and NDR status= 2 or 3 (N=21).

**Supplementary Figure 2.**
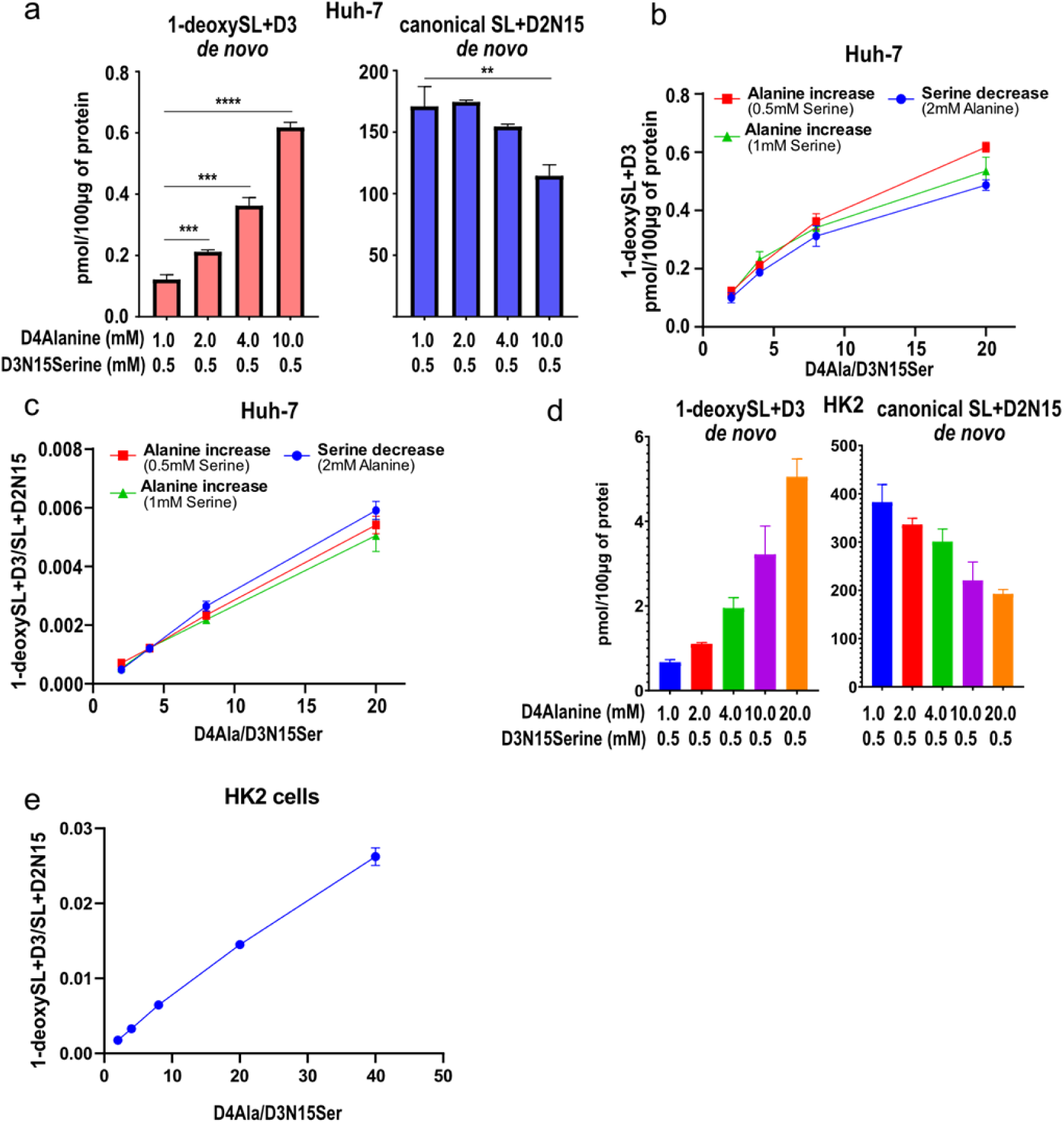
Alanine-to-Serine ratio dictates SPT activity and 1-deoxySL formation. **a** Total levels of de novo synthesized 1-deoxySL+D3 and canonical SL+D2N15 after increasing d4-L-Alanine (D4Alanine) with stable concentration of d3-15N-L-Serine (D3N15Serine) in Huh7 cells. **b** Total de novo formed 1-deoxySL+D3 amounts and **c** 1-deoxySL+D3 to SL+D2N15 ratio (1-deoxySL+D3/SL+D3) in Huh7 cells as a response to increasing D4Alanine to D3N15Serine ratio (D4Ala/D3N15Ser). Red and green represent variable L-ala against constant L-ser concentrations (0.5mM and 1mM D3N15Ser, respectively). Blue represents decreasing L-ser levels at constant L-ala concentration (2mM D4Alanine). **d** Total levels of de novo synthesized 1-deoxySL+D3 and canonical SL+D2N15 after increasing d4-L-Alanine (D4Alanine) with stable concentration of d3-15N-L-Serine (D3N15Serine) in HK2 cells**. e** 1-deoxySL+D3/SL+D3 in HK2 cells as a response to increasing D4Ala/D3N15Ser. All data are represented as mean ± SD. P-values indicate the significance of difference between the groups (two-way Student’s unpaired t-test); *=p<0.05, **=p<0.01, ***=p<0.001, ****=p<0.0001

### Increased 1-deoxySphingolipids Originate from the Liver

Having established an increased ala/ser ratio as the primary driver of 1-deoxySL formation in T2D, we next determined the source of these lipids in T2D skin. Specifically, we were interested whether 1-deoxySL are synthesized locally in skin or derived from the systemic circulation.

Skin biopsies amino acids levels revealed multiple changes in T2D patients, including alanine and serine but also glutamate, aspartate, proline, threonine, tyrosine, and branched-chain amino acids (BCAA: isoleucine, leucine, valine) (Fig. 3a). However, both serine and alanine levels were increased, leaving the ala/ser ratio unchanged (Fig. 3b). This suggests that 1-deoxySL are likely not formed directly in T2D skin but rather accumulate in response to increased circulating levels. In line with this hypothesis, we observed a significant linear correlation between plasma and skin 1-deoxySL/SL levels (Fig. 3c).

**Figure 3.**
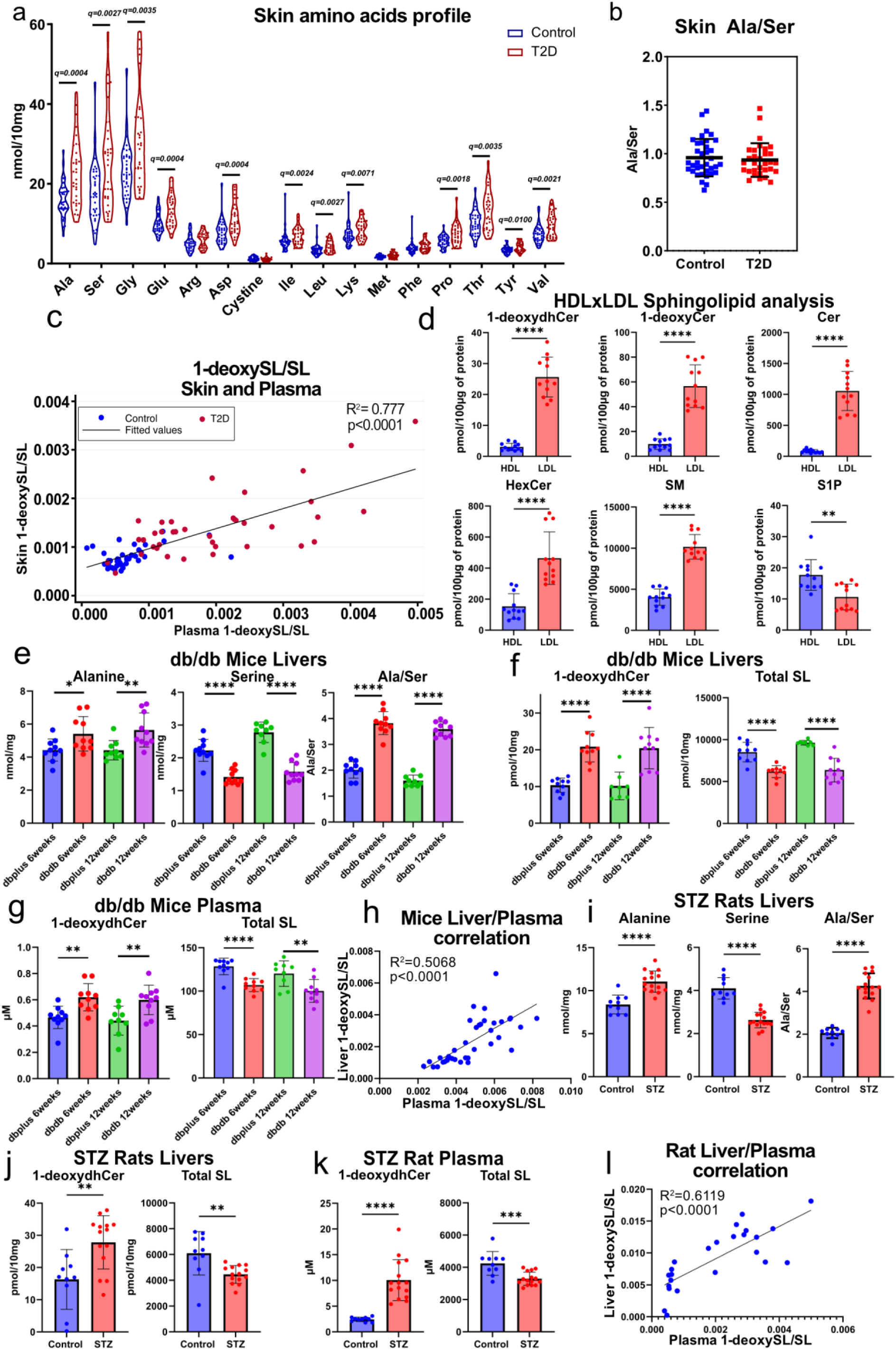
Increased 1-deoxySphingolipids are of the liver origin. **a** Violin plots showing skin amino acid levels in T2D (N=38) and healthy controls (N=39). Glycine (Gly), Glutamate (Glu), Arginine (Arg), Aspartate (Asp), Isoleucine (Ile), Leucine (Leu), Lysine (Lys), Methionine (Met), Phenylalanine (Phe), Proline (Pro), Threonine (Thr), Tyrosine (Tyr), Valine (Val) were measured using targeted LC-MS/MS metabolomics. Statistical significance was tested using Mann-Whitney and corrected for multiple testing using a two stages step-up method (Benjamini, Krieger and Yekutieli) **b** L-Alanine to L-Serine ratio (ala/ser) in skin tissue of T2D (N=38) compared to healthy controls (N=39). p reflects the significance of the group comparisons (Student’s unpaired t-test, ****=p<0.0001) **c** Correlation of plasma 1-deoxySL/SL and skin 1-deoxySL/SL. All correlations are represented as Pearson Correlation coefficient calculated from the whole cohort (N=77, red T2D, blue Controls). **d** Sphingolipidomics analysis of human plasma High density (HDL) and Low density lipoproteins (LDL). Lipoproteins were isolated from healthy donors and analyzed using untargeted LC- MS/MS lipidomics as described. **e** Amino acid and **f** Sphingolipidomics analysis of 6 and 12 weeks old, db/db (dbdb, N=10 and N=10, respectively) or db/+ (dbplus, N=10 and N=9, respectively) mouse liver tissue samples **g** Sphingolipidomics analysis of 6 and 12 weeks old, db/db (dbdb) or db/+ (dbplus) liver tissues and corresponding mouse plasma samples. **h** Correlation of liver and plasma 1-deoxySL/SL in dbdb mice. **i** Amino acid and **j** Sphingolipidomics analysis of 6 weeks old streptozotocin (STZ, N=15) or Vehicle (N=10) treated rat liver samples. **k** Sphingolipidomics analysis of 6 weeks old streptozotocin (STZ, N=15) or Vehicle (N=10) treated rat plasma samples. **l** Correlation of 1-deoxySL/SL in liver and plasma of the STZ rat vs control.

Lipids, including SL are transported in plasma mostly as lipoproteins^29^. We therefore analyzed the sphingolipid composition in the two major lipoprotein fractions, HDL and LDL from 11 healthy volunteers. We found 1-deoxySL as well as a majority of canonical sphingolipids predominantly present in LDL, while sphingosine-1-phosphate was more abundant in HDL (Fig. 3d). LDL particles – predominantly synthesized in the liver^29^– appear to be the primary carriers of 1-deoxySL, suggesting that hepatic production precedes subsequent dermal absorption.

To validate our hypothesis using accessible liver samples, we analyzed the gold standard mouse model of obesity and T2D, the db/db mouse model, which develops neuropathic-like disease^23^.

We therefore compared the lipid and amino acid profiles in liver tissues from db/db and db/+ mice at 6 and 12 weeks of age. The liver of db/db mice showed a significantly increased ala/ser ratio because of increased L-ala and reduced L-ser levels (Fig. 3e). This was paralleled with increased 1-deoxySL and reduced canonical SL levels in the db/db livers (Fig. 3f).

This hepatic lipid imbalance was also reflected in plasma. Paired plasma samples confirmed a concordant change in the 1-deoxySL/SL ratio as it was seen in liver tissues (Fig. 3g). This was further supported by the strong correlation between the hepatic and plasma 1-deoxySL/SL ratio (Fig. 3h)

To further validate our findings in a second animal model,we used an streptozotocin (STZ)-induced diabetic rat model. The STZ-treated rats also showed a significantly elevated hepatic ala/ser ratio, associated with increased L-ala and decreased L-ser levels (Fig. 3i). Consistently, lipidomics revealed increased 1-deoxySL and reduced canonical SL in liver (Fig. 3j). Again, this was mirrored in the plasma lipid profile (Fig. 3k) resulting in a strong correlation between the hepatic and plasma 1-deoxySL/SL ratio (Fig. 3l).

In summary, our results indicate that an increase in the hepatic ala/ser ratio drives hepatic 1-deoxySL formation, which are transported via LDL to peripheral tissues, including the skin. This suggests the liver as the primary source for 1-deoxySL and provides a mechanistic link between hepatic amino acid / lipid metabolism and DPN pathogenesis.

### UC13Glucose Flux Method to Investigate 1-deoxySphingolipids synthesis

L-ala is synthesized from pyruvate, while L-ser derives from 3-phosphoglycerate (3-PG); both are therefore downstream products of glycolysis. To quantitatively monitor the metabolism of glucose-derived amino acids into sphingolipid biosynthesis, we developed a UC13Glucose based metabolic flux assay to trace UC13Glucose-derived canonical and 1-deoxySL with high specificity (Fig. 4a,b).

**Figure 4.**
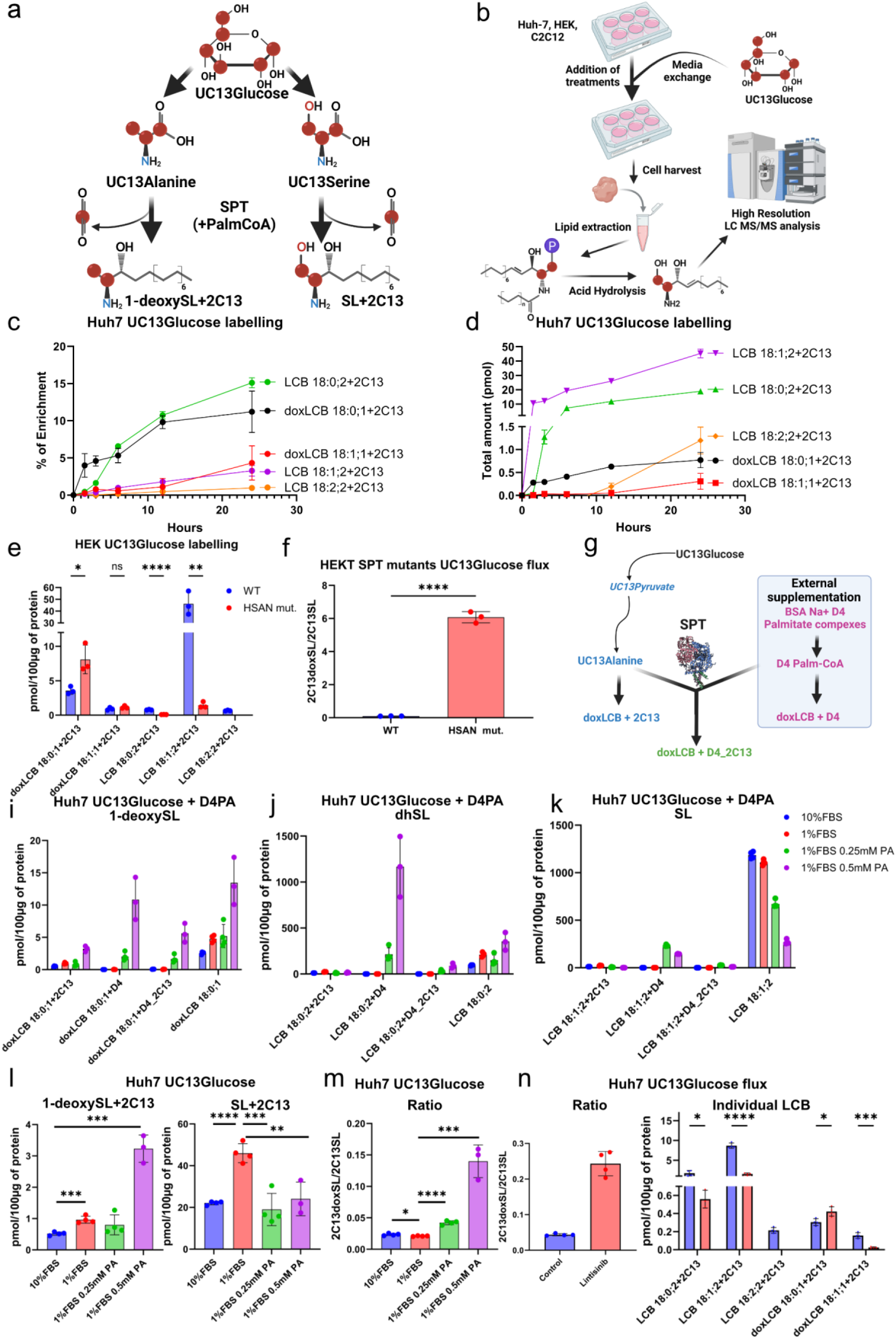
Development of a UC13Glucose Flux Method to Investigate 1-deoxySphingolipids synthesis. **a** Schematic overview of the UC13Glucose to Sphingolipids metabolic tracing assay. Glycolysis derived UC13Alanine and UC13Serine are incorporated into 1-deoxySL (1-deoxySL+2C13) and canonical SL (SL+2C13) via the SPT mediated condensation reaction **b** Optimized stable isotope labelling protocol combining UC13Glucose supplementation with an untargeted high resolution LC- MS/MS analysis after releasing the long-chain base by acid hydrolysis. Detailed procedure of the LC- MS/MS method, used internal standards and isotopic correction are indicated in the methods. **c and d** Kinetics of UC13Glucose based SL labelling in Huh7 cells. Long chain bases were analysed in relative (%) and **d** total (pmol) 2C13 enrichment after UC13Glucose supplementation. In **c** and **d** the data were normalised by the subtraction of non labelled long chain bases from Huh7 cells treated with unlabelled Glucose at the same conditions. Incorporation increased over time, peaking at 24 hours. Consequently subsequent analyses were performed at a single 24 hours time-point. **e** Total 1-deoxySL+2C13 and canonical SL+2C13 in HEK293 cells expressing either SPTLC1^WT^ (WT) or HSAN1 associated SPTLC1^C133W^ mutant SPT (HSAN mut.) after UC13Glucose supplementation. **f** The high 2C13doxSL/2C13SL ratio indicated a strongly increased 1-deoxySL formation from UC13Glucose in HSAN mut. cells. **g** Schematic of multiplexed metabolic flux assay combining UC13Glucose and D4Palmitic acid (D4PA). D4PA incorporates into 3 distinct pools of newly synthesized long-chain bases: 1.) D4 PA derived (+D4) 2.) UC13Glucose only derived (+2C13) and 3.) dual UC13Glucose and D4 PA derived (+D4_2C13) long chain bases. **i,j,k** Absolute quantification of all labelled and unlabelled 1-deoxySL dihydroSL (dhSL), and SL subpopulations demonstrating distinct metabolic regulations and confirming 2C13Glucose derived amino acids as the origin of the de-novo formed LCBs **l** Total levels UC13Glucose derived 1-deoxySL (1-deoxySL+2C13) and canonical SL (SL+2C13) and **m** 2C13doxSL/2C13SL a Huh7 cells after PA induced insulin resistance. Cells were treated with BSA Na+PA complexes in FBS deprived conditions for 3 days to induce lipotoxicity. **n** 2C13doxSL/2C13SL and total levels of UC13Glucose long chain bases in Huh7 cells treated with the insulin receptor antagonist Lintisinib. Data are represented as mean ± SD. P-values indicate the significance of difference between the groups (two-way Student’s unpaired t-test); *=p<0.05, **=p<0.01, ***=p<0.001, ****=p<0.0001

As the SPT reaction leads to a release of the C1 carbon of the utilized amino acids ^30^, the labeling with UC13Glucose will form an M+2-enriched sphingoid bases. To establish the kinetics, we analyzed the incorporation of UC13Glucose into canonical and 1-deoxySL by supplementing Huh7 cells with UC13Glucose and harvesting at multiple time points (1.5, 3, 6, 12, and 24 hours). 2C13 enrichment was detected for both, canonical SL and 1-deoxySL in a time-dependent manner, with the highest total enrichment occurring after 24 hours (Fig. 4c,d). This time point was used for the subsequent tracing assays.

The UC13Glucose flux assay was first validated in SPTLC1^C133W^ mutant expressing cells, as described earlier (Fig. 2a,e). SPTLC1^C133W^ expressing cells exhibited significantly increased UC13Glucose-derived 1-deoxySL and little UC13Glucose-derived canonical SL (Fig. 4e) thereby markedly increasing the UC13-Glucose-derived 2C13doxSL/2C13SL ratio (Fig. 4f). This confirms a glucose dependent sphingolipid formation in the cells.

To further refine our method and to ensure that 2C13 incorporation into sphingolipids really originates from UC13Glucose-derived L-ala and L-ser, we further combined the UC13Glucose tracing with teh additon of a stable isotope-labeled D4-palmitic acid (D4-PA) as a second substrate. This multiplexed metabolic flux approach allowed us to follow the formation of three distinct subpopulations of newly synthesized sphingolipids 1.) D4PA-derived LCBs (D4-LCBs), 2.) UC13Glucose-derived LCBs (2C13-LCBs) and 3.) LCBs derived from both UC13Glucose and D4-PA (D4_2C13-LCBs) (Fig. 4g). We successfully detected and quantified all three labeled and unlabeled 1-deoxySL, dihydroSL and SL populations (Fig. 4i,j,k). Importantly, the presence of D4_2C13-LCBs confirmed that the 2C13 enrichment in the de-novo formed SL originated specifically from UC13Glucose-derived amino acids.

To test the influence of metabolic stress on 1-deoxySL formation, we applied this method in an T2D in vitro model reflecting palmitic acid (PA)-induced insulin resistance, since saturated fatty acids are able to induce insulin resistance^31^. We observed a PA and dose-dependent increase in UC13Glucose derived 1-deoxySL paralleled with a decrease in canonical SL and an increased 2C13doxSL/2C13SL ratio, mirroring our findings in T2D patients.

Since insulin resistance (IR) has been linked to 1-deoxySL accumulation previously^23^, we investigated the connection between IR and changes in the UC13Glucose to SL flux using the IR/IGF1R antagonist Linsitinib^32^. With Linsitinib, the 2C13doxSL/2C13SL ratio was significantly increased which was predominantly related to a significant decrease in SL+2C13 at a modest increase in 1-deoxySL+2C13 (Fig. 4n).

Altogether, we introduce a novel UC13Glucose flux assay that, for the first time, enables precise quantification of glucose incorporation into canonical SL and atypical 1-deoxySL, providing a new tool to investigate previously uncharacterized route of sphingolipid biosynthesis.

### Proteasome-GLS-ALAT-SPT axis regulates formation of 1-deoxySphingolipids

As ala/ser ratio is increased and drives 1-deoxySL formation in T2D, we sought to search for candidate regulators of the metabolic changes underlying the imbalance of these two AAs. The alanine aminotransferase (ALAT) is a central enzyme in the alanine biosynthesis by generating alanine from pyruvate and glutamate. Therefore, we were interested in investigation whether ALAT activity contributes to the increased ala/ser ratio and 1-deoxySL formation in T2D (Fig. 5a).

**Figure 5.**
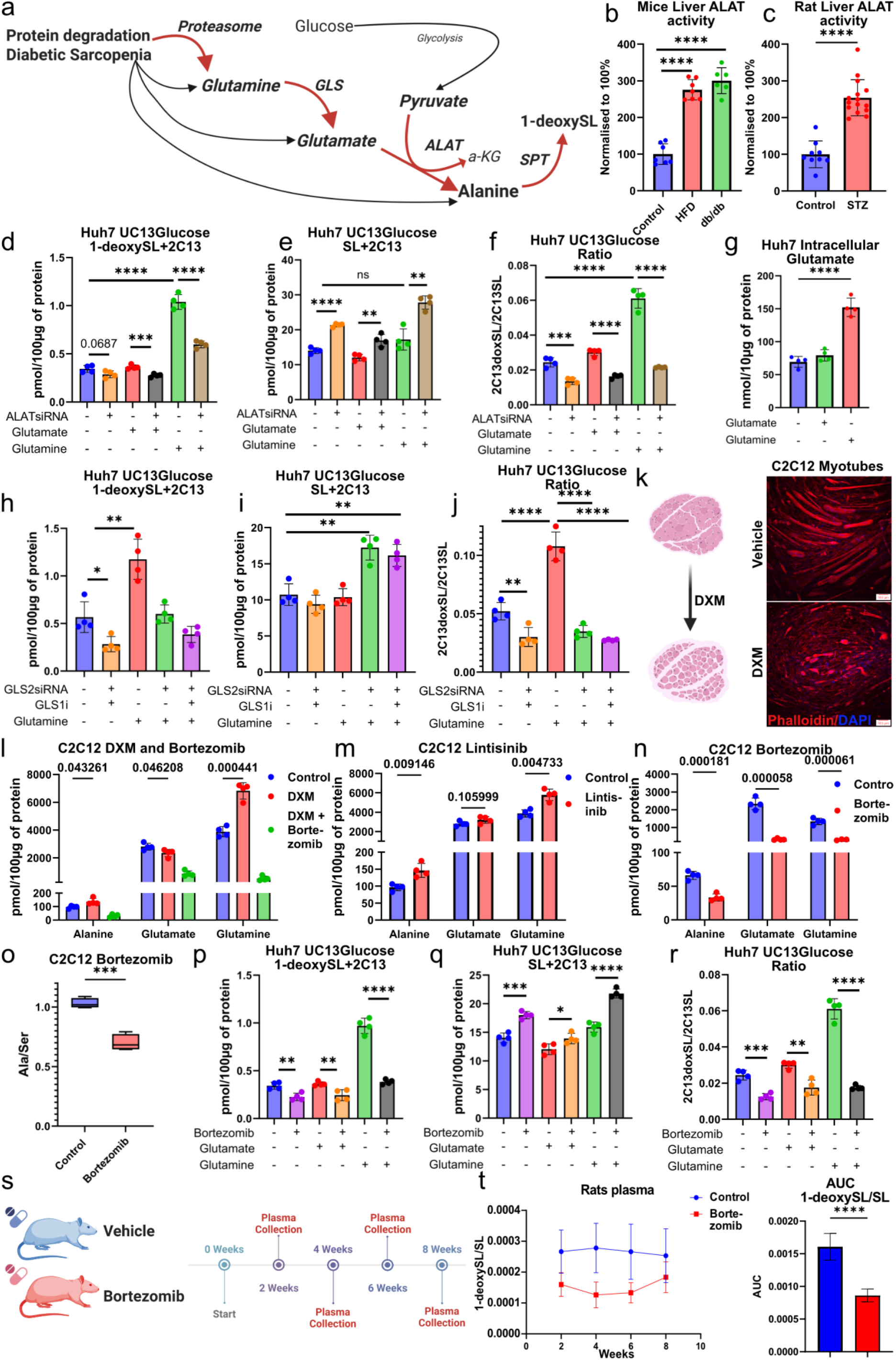
Proteasome-GLS-ALAT-SPT axis regulates formation of 1-deoxySphingolipids. **a** Schematic overview of Proteasome-Glutaminase(GLS)-Alanine aminotransferase (ALAT)-SPT axis mediated 1-deoxySL synthesis. **b,c** ALAT activity is significantly increased in livers of several diabetic rodent models (db/db, high fat diet (HFD)-fed mice and STZ treated diabetic rats) **d-f** siRNA mediated ALAT knockdown in Huh7 cells followed by UC13Glucose to SL tracing complemented with Glutamine (10mM) or Glutamate (10mM) supplementation. Total 1-deoxySL+2C13 (**d**), Total canonical SL+2C13 (**e**) and 2C13doxSL/2C13SL (**f**) were measured and calculated using high resolution LC- MS/MS, normalised to ISTD and total protein and isotopically corrected as indicated in the methods sections. **g** Total intracellular Glutamate levels in Huh7 cells after Vehicle, Glutamate (10mM) or Glutamine (10mM) treatment (24h). Amino acids were measured using targeted LC-MS/MS metabolomics **h-j** Inhibition or siRNA mediated knockdown of Glutaminases (GLS1/2) in Huh7 followed by UC13Glucose to SL tracing complemented with Glutamine (10mM) supplementation. Total 1-deoxySL+2C13 (**h**), Total canonical SL+2C13 (**i**) and 2C13doxSL/2C13SL (**j**). GLS1/2 are responsible for Glutamine to Glutamate conversion. **k** Dexamethasone (DXM)-induced diabetic sarcopenia model in differentiated C2C12 myotubes, leading to morphological degradation of myotubes as imaged by fluorescence microscopy. Scale bar=100µm. **l** Intracellular concentrations of Alanine, Glutamate and Glutamine in differentiated C2C12 myotubes after DXM treatment or combined DXM + Bortezomib treatment. **m** Intracellular concentrations of Alanine, Glutamate and Glutamine in differentiated C2C12 myotubes after insulin receptor inhibition by Lintisinib. **n** Intracellular concentrations of Alanine, Glutamate and Glutamine and **o** ala/ser in differentiated C2C12 myotubes after Bortezomib treatment. **p-r** Proteasome inhibition by Bortezomib treatment in Huh7 cells followed by UC13Glucose to SL tracing complemented with Glutamine (10mM) or Glutamate (10mM). Total 1-deoxySL+2C13 (**p**), Total canonical SL+2C13 (**q**) and 2C13doxSL/2C13SL (**r**). **s** Timeline of the Bortezomib experiment in rats. Plasma was collected at 2, 4, 6, 8 weeks of treatment. **t** Timeline of 1-deoxySL/SL in Plasma and AUC of Bortezomib compared to Vehicle treated rats. Total free long chain bases were determined after hydrolysis as specified in the methods section. All data are represented as mean ± SD. P-values indicate the significance of difference between the groups (two-way Student’s unpaired t-test); *=p<0.05, **=p<0.01, ***=p<0.001, ****=p<0.0001

Crucially, ALAT activity was significantly elevated in hepatic tissues of db/db and high-fat diet (HFD)-fed mice (Fig. 5b) as well as in STZ-induced diabetic rats (Fig. 5c) suggesting association between obesity, T2D and high ALAT activity.

To determine whether ALAT directly contributes to 1-deoxySL synthesis, we performed siRNA-mediated ALAT knockdowns (KD) in Huh7 cells, coupled with UC13Glucose flux analysis. ALAT KD significantly reduced UC13Glucose-derived 1-deoxySL formation while increasing canonical SL levels, leading to a marked decrease in the 2C13doxSL/2C13SL ratio (Fig. 5d,e,f). This demonstrates that ALAT activity is a metabolic determinant of 1-deoxySL synthesis.

Given that ALAT reversibly converts pyruvate to alanine while converting glutamate to α-ketoglutarate (α-KG), we hypothesized that glutamine metabolism (a precursor of glutamate) may influence ALAT-driven 1-deoxySL synthesis. We first evaluated whether supplementation with glutamine or glutamate could increase intracellular glutamate levels. Only glutamine supplementation increased intracellular glutamate concentrations (Fig. 5g), resulting in a marked increase in U13C-glucose-derived 1-deoxySL and an elevated 2C13doxSL/2C13SL ratio, as canonical SL levels remained unchanged (Fig. 5d, e, f). The increase was abolished in ALAT silenced cells, confirming that glutamine drives 1-deoxySL formation through ALAT (Fig. 5f).

As glutaminase (GLS) hydrolyzes glutamine into glutamate, we tested whether GLS activity contributes to 1-deoxySL biosynthesis. GLS1 (kidney-type) was inhibited using Telaglenastat (GLS1i), while GLS2 (liver-type) was silenced via siRNA knockdown. Both interventions strongly suppressed UC13Glucose-derived 1-deoxySL formation without affecting canonical SL synthesis, resulting in a significantly reduced 2C13doxSL/2C13SL ratio (Fig. 5h,i,j) independent of glutamine supplementation.

To demonstrate that glutamine serves directly as a nitrogen donor for both canonical SL and 1-deoxySL synthesis, we also applied a stable isotope tracing method using 15N (Amine)-Glutamine (15NGlutamine) (Extended Fig. 5a). Huh7 cells were treated with either vehicle, unlabeled glutamine, 15N-Glutamine, or 15N-Glutamine in combination with Myriocin, a selective SPT inhibitor. Following extraction and acid hydrolysis of long-chain bases, +15N incorporation into sphingolipids was quantified by high-resolution LC-MS/MS (Extended Fig. 5b).

15N-Glutamine, but not unlabeled glutamine, resulted in significant enrichment of +15N in both 1-deoxySL (1-deoxySL+15N) and canonical SL (SL+15N), while Myriocin treatment abolished this incorporation, confirming that 15N-glutamine is utilized by SPT for sphingolipid synthesis (Extended Fig. 5 c,d). These data provide direct evidence that nitrogen derived from glutamine contributes to both canonical and atypical sphingolipid de-novo biosynthesis.

While the role of ALAT in converting glutamate to alanine has been established, the contribution of PSAT, a glutamate-dependent aminotransferase involved in de novo serine biosynthesis, had not been investigated. Since PSAT and ALAT compete for the same substrate (glutamate), we hypothesized that PSAT activity might antagonize ALAT-mediated 1-deoxySL synthesis by channeling glutamate toward serine production and vice versa. To test this, we performed siRNA-mediated PSAT knockdown in Huh7 cells and assessed UC13C-glucose-derived sphingolipid flux. PSAT silencing significantly increased 1-deoxySL+2C13 levels and decreased canonical SL+2C13 levels, resulting in a markedly elevated 2C13doxSL/2C13SL ratio (Extended Fig. 5e–g). This indicated that reduced PSAT activity diverts glutamate away from serine synthesis and enhances alanine-mediated 1-deoxySL production.

Progressive loss of muscle mass and strength, is a common but often under recognized complication in T2D, contributing to increased frailty, reduced mobility, and poorer metabolic outcomes. Given that skeletal muscle is the primary site of glutamine production, we hypothesized that perhaps increased muscle protein degradation may contribute to elevated glutamine levels in T2D. To investigate a putative link to diabetic sarcopenia we used differentiated C2C12 myotubes as a model by inducing diabetic sarcopenia in presence of high-dose dexamethasone (DXM). DXM treatment caused visible myotube degradation (Fig. 5k) and revealed increased glutamine without seeing a major change in glutamate, and alanine levels in the atrophic myotubes.

To further test whether glutamine-driven 1-deoxySL formation is associated with increased protein degradation, we inhibited proteasome activity using the proteasome inhibitor Bortezomib (BZM). BZM treated C2C12 cells had significantly reduced intracellular glutamine, glutamate, and alanine levels (Fig. 5l) which was associated with a decreased ala/ser ratio (Fig. 5n,o). Next to see whether muscled derived Glutamine formation is mediated via IR we used Lintisinib in differentiated C2C12 myotubes which increased Glutamine and Alanine (Fig. 5m).

Consequently, proteasome inhibition by Bortezomib in Huh7 cells decreased UC13Glucose 1-deoxySL formation and increased canonical SL formation strongly decreasing 2C13doxSL/2C13SL with or without high Glutamine conditions (Fig. 5p,q,r).

The results from the cell models were also confirmed in Bortezomib treated rats. Bortezomib treated animals showed reduced plasma 1-deoxySL levels and a decreased 1-deoxySL/SL over the whole study period of 8 weeks (Fig. 5s,t), confirming that proteasome-mediated glutamine metabolism is associated with 1-deoxySL synthesis. Altogether, these data provide evidence for the Proteasome-GLS1/2-ALAT-SPT axis as the key mediator of 1-deoxySL synthesis in T2D.

**Extended Figure 5.**
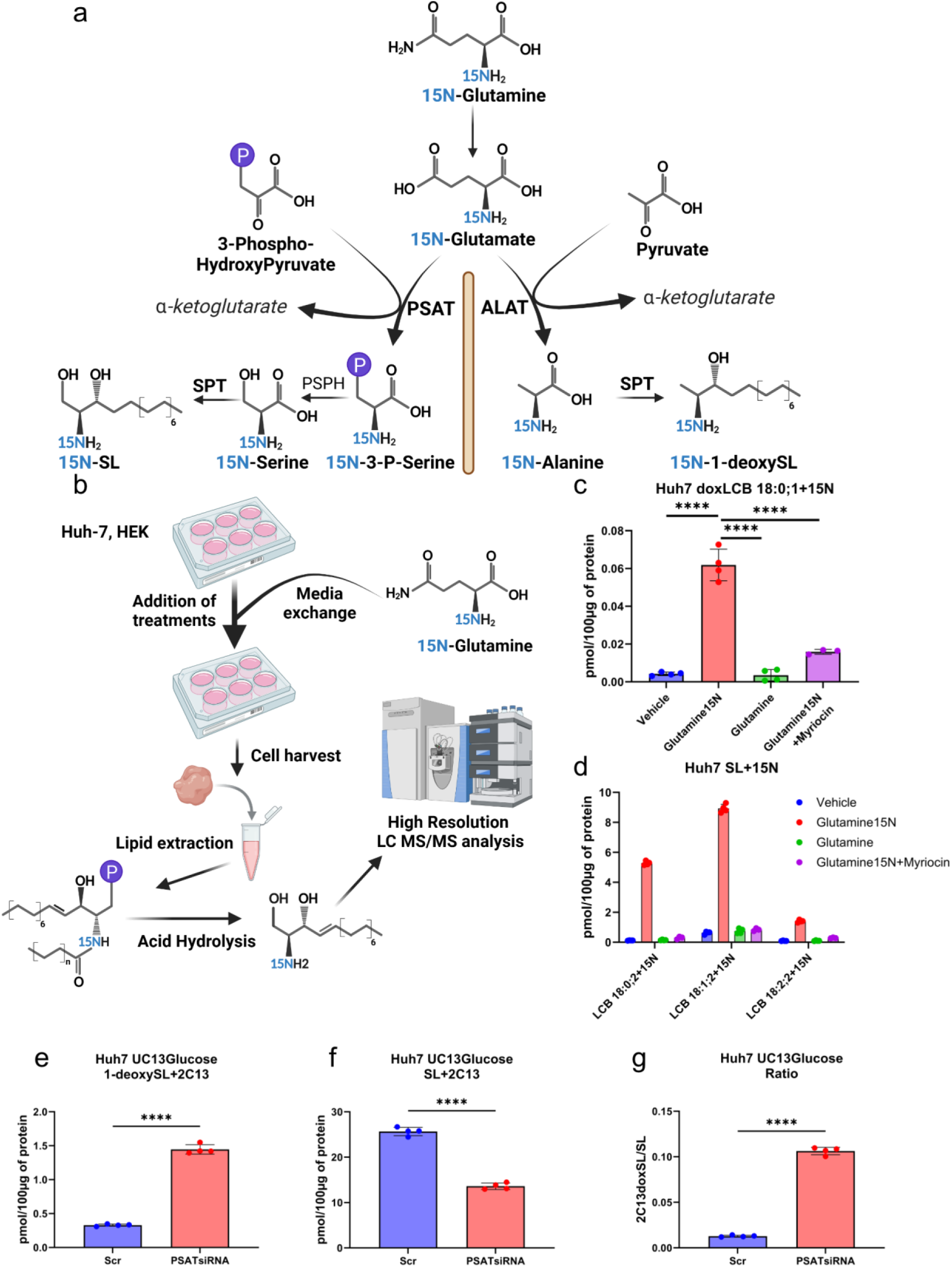
15NGlutamine to SL flux assay. **a** Scheme illustrating 15N-(Amine)Glutamine to SL flux. N15Glutamine is intracellularly converted into 15N-(Amine)Glutamate which is a substrate of the Phosphoserine amino transferase (PSAT) converting 3-Phospho-HydroxyPyruvate into a 15N-3-Phospho-Serine (15N-3-P-Serine). 15N-Glutamine derived 15N-Serine is a substrate of SPT which utilizes 15N-Serine in a canonical SL synthesis. Therefore, 15N-Glutamine derived SL will have an enrichment of +15N. On the other hand, 15N-Glutamate is also a substrate of Alanine amino transferase (ALAT) which converts Pyruvate into a 15N-Alanine. This way, 15N-Glutamine derived 15N-Alanine is utilized by SPT in formingl 1-deoxySL. Therefore, 15N-Glutamine derived 1-deoxySL will have an enrichment of +15N. **b** Optimized stable isotope labelling protocol combining 15N-Glutamine supplementation with lipid extraction and acid hydrolysis and high resolution untargeted long-chain base LC-MS/MS analysis. The experimental procedures, internal standards, LC-MS/MS method and isotopic correction are described in the methods section. **c** Total 1-deoxySL+15N in Huh7 cells after Vehicle, 15NGlutamine, Glutamine, or 15NGlutamine + Myriocin (SPT inhibitor) treatment. **d** Total canonical SL+15N in Huh cells 7 Vehicle, 15NGlutamine, Glutamine, or 15NGlutamine + Myriocin (SPT inhibitor) treatment. **e-g** siRNA mediated PSAT knockdown in Huh7 cells followed by UC13Glucose to SL tracing reflected by (**e**) total 1-deoxySL+2C13, (**f**) total canonical SL+2C13 and (**g**) 2C13doxSL/2C13SL . Total free long chain bases were determined after hydrolysis as indicated in the methods sections. All data are represented as mean ± SD. P-values indicate the significance of difference between the groups (two-way Student’s unpaired t-test); *=p<0.05, **=p<0.01, ***=p<0.001, ****=p<0.0001

### L-serine metabolism modulates the formation of 1-deoxySphingolipids

As ala/ser ratio is the key determinant of 1-deoxySL formation in T2D, we were also interested about the contribution of the serine metabolism on 1-deoxySL formation (Fig. 6a). We therefore combined our UC13Glucose flux assay with the supplementation of stable isotope-labelled amino acids that allowed a systematic investigation of serine homeostasis and its regulatory role in 1-deoxySL biosynthesis.

**Figure 6.**
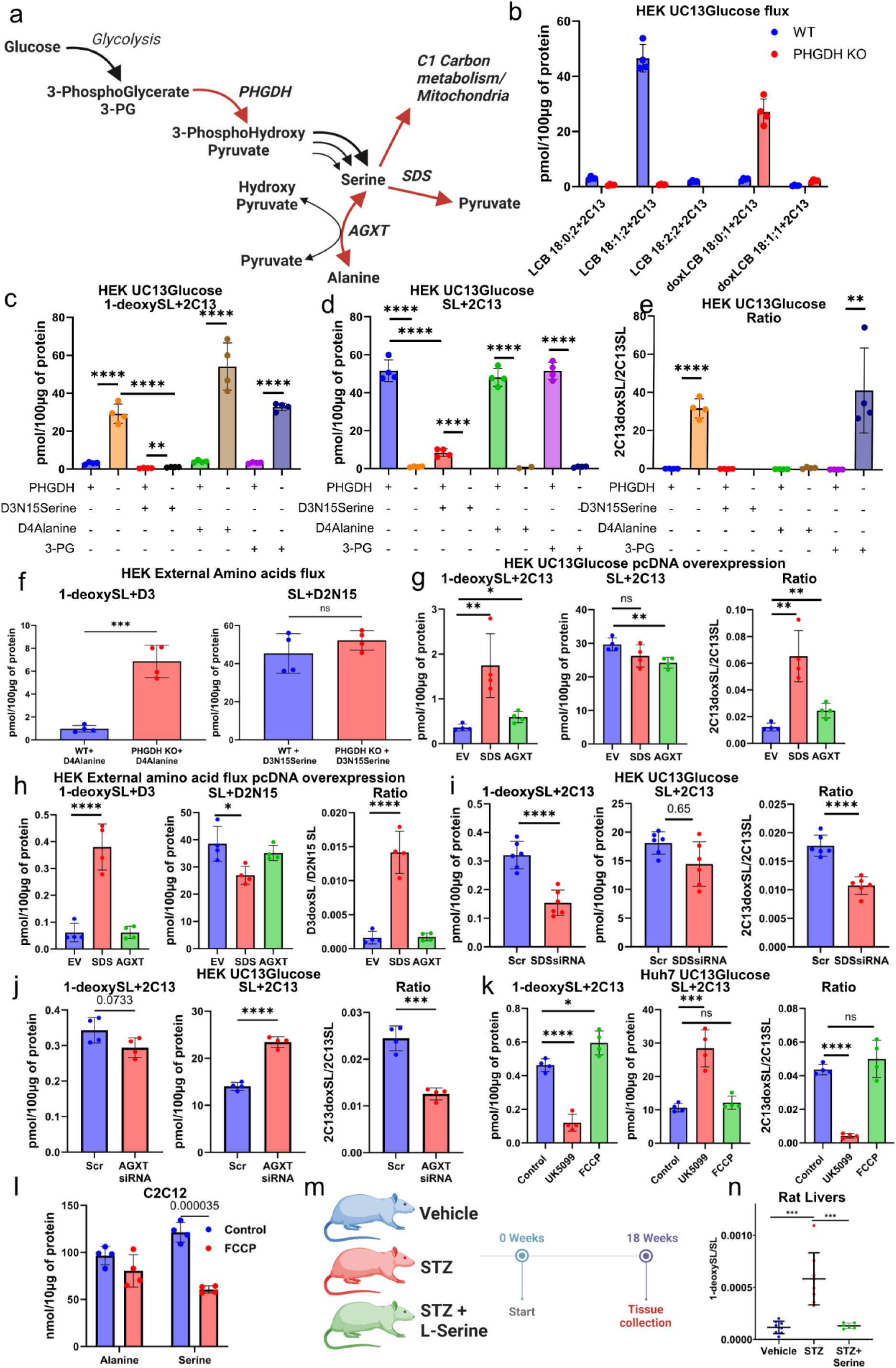
L-serine homeostasis regulates formation of 1-deoxySphingolipids. **a** Schematic overview of serine biosynthetic and catabolic pathways. **b** UC13Glucose to SL tracing in PHGDH knockout (PHGDH KO) HEK293 cells. **c-e** UC13Glucose to SL tracing complemented with D3N15Serine, D4Alanine or 3-phosphoglycerate (3PG) supplementation. Total 1-deoxySL+2C13 (c), Total canonical SL+2C13 (d), 2C13doxSL/2C13SL (e), 1-deoxySL+D3 (supplemented d4Alanine derived, f), SL+D2N15 (supplemented D3N15Serine derived, **f-h** Overexpression of serine-degrading enzymes, Serine dehydratase (SDS) and alanine-glyoxylate aminotransferase (AGXT) or empty vector (EV) in HEK293 cells, combined with UC13Glucose to SL tracing. h Influence of SDS, AGXT or EV overexpression in HEK293 cells on D4Alanine derived 1-deoxySL+D3 and D3N15Serine derived canonical SL+D2N15 as well as the ratio between 1-deoxySL/SL. **i** siRNA mediated knockdown of SDS in HEK293 cells coupled with UC13Glucose to SL tracing. **j** siRNA mediated knockdown of AGXT in HEK293 cells coupled with UC13Glucose to SL tracing. **k** Effect of inhibiting the Mitochondrial Pyruvate carrier protein (MPC) by UK5099 or by uncoupling OxPhos with FCCP on the UC13Glucose to SL flux in Huh7 cells. **l** Influence of FCCP treatment on intracellular L-ala and L-ser concentrations in differentiated C2C12 myotubes. All lipids and metabolites were quantified using high resolution LC-MS/MS as indicated in the methods sections. **m** Scheme illustrating experiment with L-Serine supplementation on STZ treated Rats. **n** Liver 1-deoxySL/SL ratio in Vehicle, STZ or STZ + L-Serine treated rats. Lipids were measured by high resolution LC-MS/MS lipidomics as described in method section. All data are represented as mean ± SD. P-values indicate the significance of difference between the groups (two-way Student’s unpaired t-test); *=p<0.05, **=p<0.01, ***=p<0.001, ****=p<0.0001

To assess the role of phosphoglycerate dehydrogenase (PHGDH), the rate-limiting enzyme in serine biosynthesis, we performed an UC13Glucose flux assay in PHGDH deficient HEK293 cells (PHGDH KO). The KO cells showed strongly reduced synthesis of UC13Glucose-derived canonical SL and increased 1-deoxySL synthesis (Fig. 6b,c,d,e).

To further investigate the effect of serine deficiency on SL formation, we supplemented PHGDH KO cells with either 3-phosphoglycerate (3-PG), the upstream precursor of serine biosynthesis, or D3N15-labeled serine (D3N15Serine). 3-PG supplementation had no effect on 1-deoxySL formation, whereas D3N15Serine supplementation completely abolished UC13Glucose-derived 1-deoxySL synthesis (Fig. 6c) in PHGDH KO cells. Importantly, d3N15Serine supplementation also suppressed formation of canonical SL+2C13, possibly by competing with endogenously formed serine+2C13 or by a serine-mediated feedback inhibition of PHGDH (Fig 6d). This confirms that serine deficiency enhances 1-deoxySL synthesis by switching to alanine as an alternative substrate, resulting in an increased 1-deoxySL/SL (Fig 6e).

To assess whether serine insufficiency increases the alanine utilisation by SPT, we supplemented PHGDH KO cells with D4-alanine and measured its incorporation into 1-deoxySL (1-deoxySL+D3). The addition of D4-alanine significantly increased D3-labeled 1-deoxySL in PHGDH KO cells, whereas supplementation of D3N15Serine had no stimulatory effect on canonical SL+D2N15 (Fig f). This again demonstrate that low serine levels, causing increased ala/ser ratio, push SPT to utilize alanine, promoting 1-deoxySL accumulation.

We also tested the contribution of the serine dehydratase (SDS, which reversibly metabolizes serine to pyruvate) and alanine-glyoxylate aminotransferase (AGXT, which reversibly metabolizes pyruvate to alanine) on 1-deoxySL formation. Overexpressing SDS and AGXT in HEK293 cells (Fig 6a) increased UC13Glucose-derived 1-deoxySL while slightly decreasing canonical SL levels, increasing the 2C13doxSL/2C13SL ratio (Fig 6g). Upon supplementation with D4-alanine and D3N15-serine, SDS-overexpressing cells showed a significant increase in the 1-deoxySL+D3/SL+D2N15 ratio, whereas AGXT overexpression did not significantly affect either sphingolipid species (Fig 6h).

To further assess whether SDS and AGXT represent viable targets for reducing 1-deoxySL levels, we also performed siRNA-mediated knockdowns of these enzymes. Interestingly, SDS knockdown strongly suppressed UC13Glucose-derived 1-deoxySL synthesis with only a minimal reduction in canonical SL levels, leading to a substantial decrease in the 2C13doxSL/2C13SL ratio (Fig 6 i). AGXT knockdown however, had a weaker effect, modestly decreasing 1-deoxySL while significantly increasing canonical SL formation and therefore strongly reducing 1-deoxySL/SL (Fig 6j).

Given the predominant hepatic expression of SDS and AGXT, we also confirmed their functional relevance using hepatocyte Huh7 cells as a model (Supplementary Fig 6 a). siRNA-mediated knockdown of SDS and AGXT in Huh7 cells, followed by UC13Glucose flux analysis, significantly reduced 1-deoxySL (1-deoxySL+2C13) in parallel with a concomitant increase in SL+2C13, resulting in a overall decrease in the 2C13doxSL/2C13SL ratio (Supplementary Fig. 6b–d). These findings support a regulatory role for SDS and AGXT in controlling sphingolipid substrate flux and suggest they may represent novel therapeutic targets to lower 1-deoxySL levels.

Serine metabolism is closely linked to mitochondrial function, particularly through mitochondrial pyruvate transport, which has been implicated in 1-deoxySL regulation^33^. To assess whether also mitochondrial dysfunction influences serine-dependent sphingolipid metabolism, we inhibited the mitochondrial pyruvate carrier (MPC) using UK5099 (MPCi) and induced mitochondrial oxidative stress using FCCP.

Inhibition of MPC strongly suppressed UC13Glucose-derived 1-deoxySL formation and increased canonical SL synthesis resulting in a decreasing 1-deoxySL/SL (Fig 6 k). Presence of FCCP had only a mild effect on both lipid species (Fig 6k). To further explore the link between mitochondrial dysfunction and serine metabolism, we also treated differentiated C2C12 myotubes with FCCP and measured intracellular amino acid levels. FCCP treatment significantly reduced intracellular serine levels, while alanine concentrations remained unchanged (Fig 6l). These findings indicate that oxidative stress-induced mitochondrial dysfunction may contribute to serine depletion, further exacerbating 1- deoxySL accumulation in T2D.

Finally, to assess whether the expression of serine biosynthetic and catabolic enzymes is altered in vivo under diabetic conditions, we examined hepatic mRNA expression levels of *Phgdh*, *Sds*, *Agxt*, *Shmt1*, and *Shmt2* in *db/db* mice at 6 and 12 weeks of age and in STZ-induced diabetic rats. *Phgdh* expression was modestly reduced in *db/db* mice at 6 weeks and unchanged at 12 weeks and in STZ-treated rats (Supplementary Fig. 6e,j). *Sds* expression showed a progressive increase in *db/db* mice but was markedly suppressed in STZ rats (Supplementary Fig. 6f,k). *Agxt* was slightly downregulated in *db/db* mice at 6 weeks and strongly reduced in STZ livers (Supplementary Fig. 6g,l).

Given the link between mitochondrial function, oxidative stress, and serine homeostasis, we also assessed expression of *Shmt1* and *Shmt2*, which catalyze the conversion of serine to glycine, a precursor for glutathione (GSH) synthesis. In theory, increased SHMT activity could deplete intracellular serine and promote alanine-mediated 1-deoxySL synthesis. *Shmt1* and *Shmt2* expression was increased in *db/db* mouse livers at 12 weeks, but both genes were significantly downregulated in STZ rats (Supplementary Fig. 6h,i,m,n).

These results suggest that transcriptional changes in hepatic serine metabolic can only partially explain the serine depletion and increased 1-deoxySL levels observed in type 2 diabetes. However, functional modulation of SDS and AGXT revealed a direct impact on sphingolipid flux, establishing these enzymes as promising targets for therapeutic intervention in 1-deoxySL-related metabolic complications.

Given that serine deficiency enhances 1-deoxySL formation, we also investigated whether L-serine supplementation could mitigate 1-deoxySL accumulation in vivo. STZ-induced diabetic rat were therefore treated with an serine enriched diet for 18 weeks (Fig 6 m).

In STZ-treated rats, the hepatic 1-deoxySL/SL ratio was elevated in the absence of treatment; however, oral L-serine supplementation restored the ratio to baseline levels (Fig. 6n). These findings offer in vivo evidence that L-serine supplementation is a straightforward, effective, and safe approach to reduce neurotoxic 1-deoxySL levels in T2D.

**Supplementary Figure 6.**
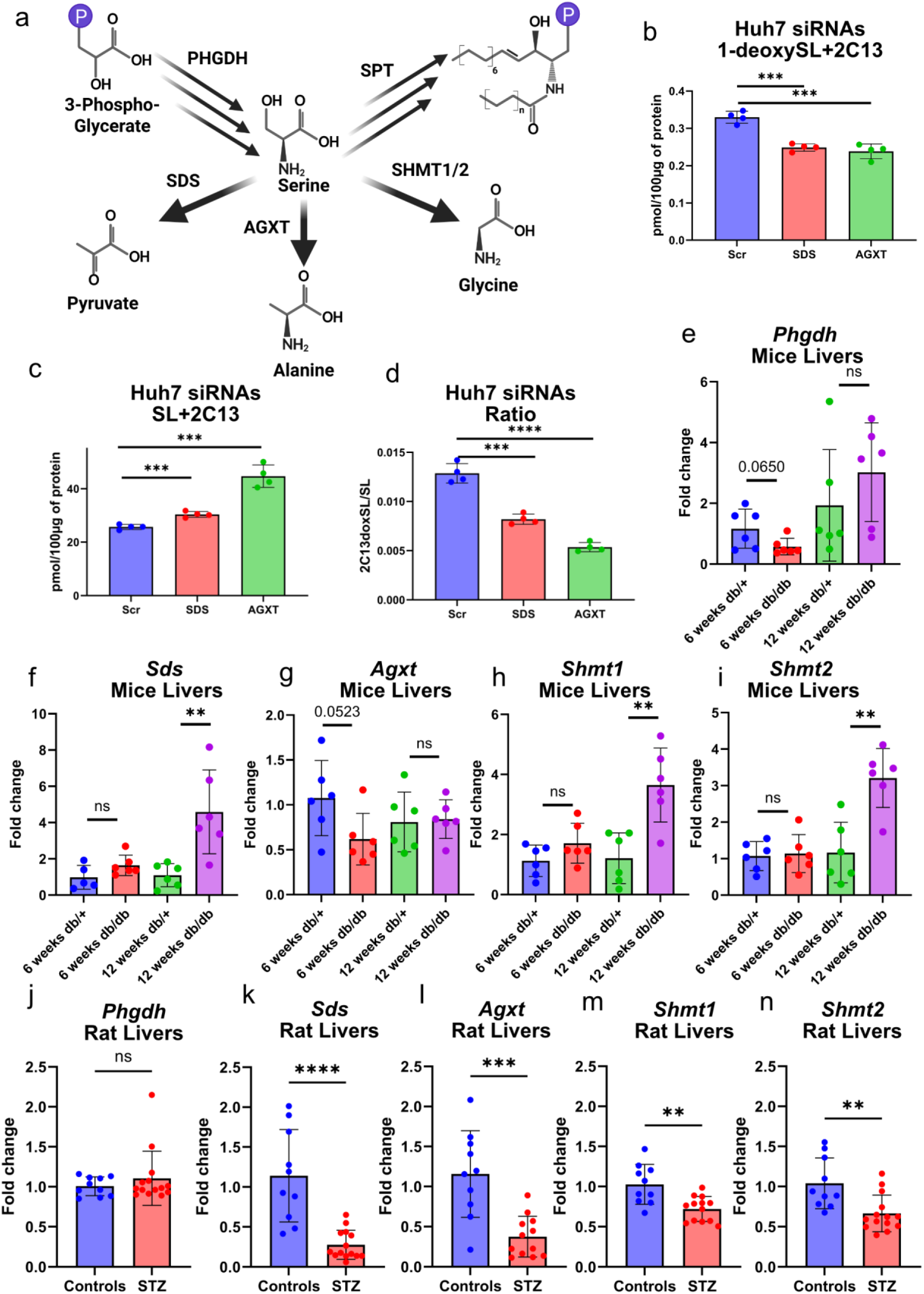
**a** L-Serine biosynthesis and catabolism and its connection to SL synthesis n respect to Phosphoglycerate dehydrogenase (PHGDH), Serine dehydratase (SDS), alanine-glyoxylate aminotransferase (AGXT) and Serine hydroxymethyl transferase 1/2 (SHMT1/2). **b-d** Total 1-deoxySL+2C13 in siRNA mediated SDS and AGXT knockdown followed by UC13Glucose to SL tracing in Huh7 cells. **(b)** total canonical SL+2C13 **(c)** 2C13doxSL/2C13SL **(d)** total free long chain bases as determined after hydrolysis as indicated in the methods sections. **e-i** Expression levels of hepatic *Phgdh* **(e)**, *Sds* **(f)**, *Agxt* **(g)**, *Shmt1* **(h)**, and *Shmt2* **(i)** in *db/db* and *db/+* mice at 6 and 12 weeks of age (N=10 for each group, except *db/+* at 12 weeks: N=9), assessed by quantitative PCR (qPCR). **j–n** Hepatic expression of *Phgdh* **(j)**, *Sds* **(k)**, *Agxt* **(l)**, *Shmt1* **(m)**, and *Shmt2* **(n)** in STZ-treated and vehicle-treated (control) rats, as determined by qPCR. All data are represented as mean ± SD. P-values indicate the significance of difference between the groups (two-way Student’s unpaired t-test); *=p<0.05, **=p<0.01, ***=p<0.001, ****=p<0.0001

## Discussion

In this study, we evaluated the metabolism of 1-deoxySL in T2D patients. Our findings indicate that 1-deoxySL are not only elevated in T2D plasma but are also significantly increased in the skin of diabetic patients, which associates with clinically incident DPN in our cohort. Moreover, we demonstrate that, irrespective of genetic alterations, metabolic changes in amino acids levels modulate 1-deoxySL biosynthesis. Together, our study provides a comprehensive metabolic framework linking an increased ala/ser ratio to increased 1-deoxySL formation in T2D and DPN.

1-deoxySL are a class of atypical, neurotoxic sphingolipids generated by SPT when the enzyme metabolizes L-alanine instead of its preferred substrate L-serine. The monogenic neuropathy HSAN1 is driven by elevated 1-deoxySL levels, resulting in peripheral sensory neuropathy due to a sustained substrate shift towards L-alanine. Notably, HSAN1 and the diabetic sensory neuropathy share a similar clinical presentation, characterized by a length-dependent and progressive sensory loss, impaired wound healing, ulcerations, and neuropathic pain. The pathological 1-deoxySL formation seen in both conditions, suggests a shared pathological mechanism.

Elevated plasma 1-deoxySL levels have also been reported in metabolic conditions including MAFLD and T2D. Since no SPT mutations are associated with MAFLD or T2D, a permanent shift in the SPT substrate specificity does not explain the increase in 1-deoxySL. Instead, we demonstrated that the increase in 1-deoxySL/SL is primarily associated with a relative increase in the ala/ser ratio and largely independent of absolute amino acid concentrations. We demonstrated a strong linear relationship between 1-deoxySL/SL and ala/ser ratios in both cellular and animal models (Fig. 2a,c).

Changes in the amino acid profiles are a well-established metabolic hallmark of T2D. Recent studies demonstrated that high-fat diet-induced insulin resistance leads to L-serine deficiency, increased 1-deoxySL levels, and DPN in experimental mouse models^23^. Under L-serine restriction, combined with obesity and insulin resistance, mice exhibit increased 1-deoxySL synthesis and develop sensory neuropathy. Notably, L-serine supplementation or pharmacological inhibition of SPT (e.g., myriocin) alleviated neurotoxic effects by reducing 1-deoxySL levels^23^. Consistent with these findings, we observed reduced L-serine levels in our T2D cohort. However, L-serine deficiency alone does not appear to be the sole driver of 1-deoxySL accumulation, as we also found significantly elevated L-alanine levels, which means that increased ala/ser ratio in T2D is caused from both sides. Using various cell culture models, we demonstrated that increased ala/ser ratio directly induces 1-deoxySL formation, contributing to DPN in T2D (Fig. 2, Fig. 3, Extended Fig. 2).

Our findings also indicate that 1-deoxySL primarily originates in the liver and is transported via LDL to peripheral tissues, including the skin. A strong correlation between plasma and skin 1-deoxySL/SL ratios supports a systemic origin (Fig. 2d). Further lipidomics analysis of plasma lipoproteins confirmed that LDL, rather than HDL, serves as the primary transporter of 1-deoxySL (Fig. 3c,d). Consistently, lipidomics and amino acid profiling of diabetic rodent models revealed a significant increase in L-alanine and a decrease in L-serine which results in an increased ala/ser ratio. Thus, the liver appears as a key source for circulating 1-deoxySL in T2D.

To further elucidate the metabolic pathways contributing to 1-deoxySL accumulation, we developed multiplexed UC13Glucose to SL flux assays. This approach allowed to metabolically trace glucose-derived amino acids (L-serine and L-alanine) into sphingolipid synthesis. Using this multiplexed method and also incorporating D4-palmitic acid (D4-PA), we identified distinct subpopulations of newly synthesized sphingolipids, confirming that 1-deoxySL formation increases under lipotoxic/insulin-resistant conditions. Notably, the inhibition of the insulin receptor further enhanced the 1-deoxySL/SL ratio by lowering canonical SL synthesis. These findings underscore the relevance of insulin signaling in SL homeostasis and establish UC13Glucose flux assays as a powerful tool for studying SL metabolism in metabolic diseases. ALAT has been connected as one of the regulators of T2D pathogenesis^34^. Increased ALAT expression has been demonstrated in patients and animal models with T2D and reducing ALAT activity prevented from muscle atrophy and hyperglycemia in vivo^34^.

Consistently, we found that liver alanine aminotransferase (ALAT) activity was significantly elevated in various diabetic rodent models while ALAT knockdown in hepatic cells resulted in a marked reduction in UC13Glucose-derived 1-deoxySL formation, confirming ALAT’s direct influence on 1- deoxySL metabolism.

Complementary, also glutamine metabolism emerged as a key upstream regulator of this pathway. Glutaminase (GLS1/2) inhibition significantly reduced UC13Glucose-derived 1-deoxySL, further linking glutamine-derived glutamate to ALAT-mediated alanine production (Fig. 5h-j).

Diabetic sarcopenia is well-established complication of T2D^35^, associated with altered activity of ubiquitin-proteasome system and proteolytic flux^36^. In this study, we demonstrate that dexamethasone-induced diabetic sarcopenia leads to elevated glutamine levels, suggesting a mechanistic link between muscle protein degradation and increased 1-deoxySL synthesis. Consistent with this, pharmacological inhibition of the proteasome by Bortezomib, reduced intracellular glutamine and alanine concentrations, and lowered hepatic 1-deoxySL production with or without Glutamine supplementation (Fig. 5n,o). Importantly, these effects also translated in vivo, as proteasome inhibition by Bortezomib in rats significantly reduced plasma 1-deoxySL levels over time (Fig. 5s,t). It should be noted that although Bortezomib is clinically associated with peripheral neuropathy, this effect is not directly attributable to proteasome inhibition, as the more selective proteasome inhibitor Carfilzomib does not induce similar neurotoxic effects^37^. In summary, our data establish a proteasome-GLS-ALAT-SPT axis as a critical regulator of 1-deoxySL formation in T2D.

Further, we demonstrated the role of L-serine homeostasis in regulating 1-deoxySL formation in more detail^23^. Deficiency in phosphoglycerate dehydrogenase (PHGDH), the rate-limiting enzyme in L-serine biosynthesis, reduced ala/ser ratio and increased 1-deoxySL formation in the UC13Glucose flux assay. This highlights the importance of the L-ser and L-ala balance in SPT substrate selection. Furthermore, PHGDH deficiency is causing peripheral sensory neuropathy as shown previously in patients and also in rodent models^38–40^. Interestingly, we found that another L-ser synthesizing enzyme, phosphoserine aminotransferase PSAT1 regulates 1-deoxySL formation, and has also been associated with the development of juvenile-onset neuropathy^41^. Moreover, we demonstrated that SDS and AGXT, two other enzymes involved in serine metabolism effectively modulate 1-deoxySL formation, making them potential targets to suppress the formation of these neurotoxic lipids.

Mitochondrial dysfunction and increased oxidative stress are one of the hallmarks in T2D^42^. We demonstrated that inhibition of mitochondrial pyruvate transport (MPCi) lowers 1-deoxySL levels while increasing canonical SL synthesis, further supporting a link between mitochondrial metabolism and serine availability. Lastly, we also showed that FCCP treatment effectively reduced L-serine levels in differentiated myotubes, which suggests that mitochondrial dysfunction and oxidative stress also influence L-serine homeostasis and contribute to changes in the ala/ser ratio in T2D.

L-serine supplementation in HSAN1 has been proven to reduce 1-deoxySL levels in clinical studies and to alleviate neuropathic symptoms^24^. We reconfirmed that oral L-serine supplementation can also effectively reduce hepatic and plasma 1-deoxySL levels in diabetic rats. This reinforces the therapeutic potential of serine-based interventions in DPN and its potential for DPN management.

In conclusion, our study provides a comprehensive metabolic framework linking an increased ala/ser ratio to increased 1-deoxySL formation in T2D and DPN. We identify the liver as a key site of 1- deoxySL production, with transport occurring primarily via LDL. Also, we established the Proteasome-GLS-ALAT-SPT axis, L-Serine biosynthesis complemented with L-Serine catabolism as major metabolic regulators of 1-deoxySL synthesis. Finally, we introduce UC13Glucose flux assays as a novel tool to investigate SL metabolism and identify new upstream targets. Lastly, we proposed that L-serine supplementation as a potential therapeutic strategy for reducing neurotoxic 1-deoxySL levels.

Further interventional studies are warranted to confirm the efficacy of L-serine in clinical settings and to explore whether similar metabolic disturbances occur in other conditions associated with muscle wasting, such as aging or cancer-related cachexia. Targeted therapies aimed at restoring amino acid homeostasis could provide novel avenues for managing diabetic neuropathy and related metabolic complications.

## Material and methods

### Study subjects

We obtained plasma and skin biopsy samples from a group of individuals with type 2 diabetes (T2D) and healthy sex- and age-matched controls who were recruited by Dr Pär Björklund at the Karolinska Institute in Stockholm, Sweden, during 2018-2019. This cohort was also used for research on the response-to-retention hypothesis of atherogenesis (ClinicalTrials.govID NCT03386097), which involved taking punch biopsies to measure lipid accumulation in the skin. The study was approved by the local ethics committee in Stockholm (Approval Number 2017-1942-31, 2018-1428-32, 2019-06474, 2021-06485-02) and was conducted in accordance with the Declaration of Helsinki. The inclusion criteria for the study at Karolinska was the diagnosis of type diabetes mellitus according to the Classification of Diabetes Mellitus 2019 (WHO). Exclusion criteria for the T2D group included thyroid disease, systemic inflammatory disease, plasma levels of haemoglobin, thyroid hormones or inflammatory markers outside of the laboratory reference range, treatment with oral glucocorticoids, pregnancy, and systemic skin disease. The exclusion criteria for the healthy controls were blood pressure over 140/90, systemic skin disease, laboratory analyses outside of laboratory reference range, pregnancy, and non-trivial disease requiring treatment, except for hypertension treated with no more than one antihypertensive drug that should not be an alpha-blocker, beta-blocker or thiazide-diuretic. T2D subjects were mainly recruited from primary care centres in Stockholm; healthy controls were also recruited from the Stockholm area. The number of T2D subjects and controls at each stage of the study and the reason for non-participation are presented in Supplementary Fig. 6. The main clinical characteristics of the sex and age-matched T2D subjects and controls are summarised in Table 1. The data used for screening was obtained from anamnesis taken by PB, from health forms, patient records, clinical examinations, and routine plasma analyses at Karolinska University Hospital Huddinge. The average disease duration was 10 years, with 18% having experienced a prior major cardiovascular event (MACE) (acute myocardial infarction n=7, stroke n=0). 55% had signs of peripheral neuropathy (NDR2 n=20, NDR3 =1), 8% had retinopathy, and 16% had nephropathy. Tow-thirds of T2D subjects were on long-term antidiabetic treatment (metformin n=27, Insulin n=12, GLP1-agonist n=3, SGLT2-inhibitor n=4, DPP4-inhibitors n=8, sulfonylurea n=2), 74% were on lipid-lowering therapy (Simvastatin n=7, Atorvastatin n=20, Rosuvastatin n=1, Ezetimib n=3). There was no significant difference in kidney function (eGFR) or level of systemic inflammation (hs-CRP) between T2D subjects and healthy controls, but the systolic blood pressure was 8 mmHg higher in T2D subjects.

**Table 1.** Demographic and clinical characteristics of participants.

|  | **C** | **T2D** |
| --- | --- | --- |
| **Males : Females (n:n)** | 24:15 | 24:14 |
| **Age (y)** | 68.7 ± 10.6 | 66.9 ± 10.4 |
| **BMI (kg/m<sup>2</sup>)** | 25.1 ± 2.6 | 28.2 ± 4.0 *** |
| **HbA1c (mmol/mol)** | 37.1 ± 3.8 | 57.2 ± 15.3 *** |
| **FBG (mM)** | 5.4 ± 0.4 | 7.7 ± 2.5 *** |
| **SBP (mmHg)** | 131 ± 14 | 139 ± 16 * |
| **eGFR(CysC)relative (mL/min/1.73m<sup>2</sup>)** | 76.4 ± 11.4 | 70.5 ± 16.3 |
| **hsCRP (mg/l)** | 1.17 ± 1.16 | 1.64 ± 2.05 |
| **Diabetes duration (y)** | - | 9.6 ± 8.8 |
| **Antidiabetic treatment (n)** | - | 32 |
| **Lipid lowering therapy (n)** | 2 | 28 |
| **Antihypertensive treatment (n)** | 8 | 30 |
| **Prior MACE (n)** | - | 7 |
| **Nephropathy (n)** | - | 6 |
| **Retinopathy (n)** | - | 3 |
| **NDR1 = no evidence of distal neuropathy (n)** | - | 17 |
| **NDR2 = signs of distal neuropathy (n)** | - | 20 |
| **NDR3 = signs of distal neuropathy + previous foot ulcers, amputation, foot deformity or other skin deformity (n)** | - | 1 |
| **NDR4 = ongoing foot ulcers regardless of neuropathy or vascular disease, or severe osteopathy or pain syndrome** | - | - |

### Collection of skin punch biopsies

Skin punch biopsies and plasma were collected at the Clinical Science Center at Karolinska University Hospital Huddinge. Participants arrived in the morning after at least 8 hours of fasting, and anthropometric measurements were registered, followed by phlebotomy. Three 4mm punch biopsies were taken from the skin on the belly, just over 5 centimeters lateral to the belly button. The subcutaneous fat was removed from the dermis and epidermis under a microscope using a knife, and the biopsies were then snap-frozen on dry ice and stored at -80°C.

### Neuropathy data

Data on neuropathy registered according to the Swedish National Diabetes Registry’s (NDRs) standard for diabetic foot assessment were obtained from patients’ records. On average, the NDR status had been assessed and recorded 118 days prior to the date of the skin biopsy (ranging from 496 before the biopsy to 366 after). Diabetic peripheral neuropathy (DPN) can be diagnosed based on clinical or electrophysiological criteria. One widely accepted definition of DPN is “the presence of symptom and/or signs of peripheral nerve dysfunction in people with diabetes after exclusion of other causes”. In Sweden, neuropathy and angiopathy in T2D patients are determined and recorded yearly according to the Swedish National Diabetes Registry’s (NDRs) standard for diabetic foot assessment. The assessment is done by a nurse specialised in diabetes and includes a) a careful inspection of the patient’s feet, b) palpation of arteria dorsalis pedis and arteria tibialis posterior, c) an interview of the patient about symptoms of tingling and/or numbness in the feet, d) assessment of pinprick sensation with monofilament 5,07-10g at the tip of digitus 1, and at the medial and lateral aspects of the ball of the foot, e) assessment of vibration perception at the medial malleolus, at the base of digitus 1, and at the tip of digitus 1 using a 128 Hz tuning fork. If there are no signs of distal neuropathy, peripheral vascular disease or other foot problems, the patient is classified as NDR1. If there are signs of distal neuropathy or peripheral vascular disease based on the absence of pulses in any of the arteries or the absence of pinprick or vibration perception at any location, the patient is classified as NDR2. Patients are classified as NDR3 if there are signs of distal neuropathy or peripheral vascular disease, plus a history of foot ulcers, amputation, foot deformity, or other skin pathology such as calluses and skin cracks. If there are ongoing foot ulcers, regardless of neuropathy or vascular disease, severe osteopathy or pain syndrome, the patient is classified as NDR4

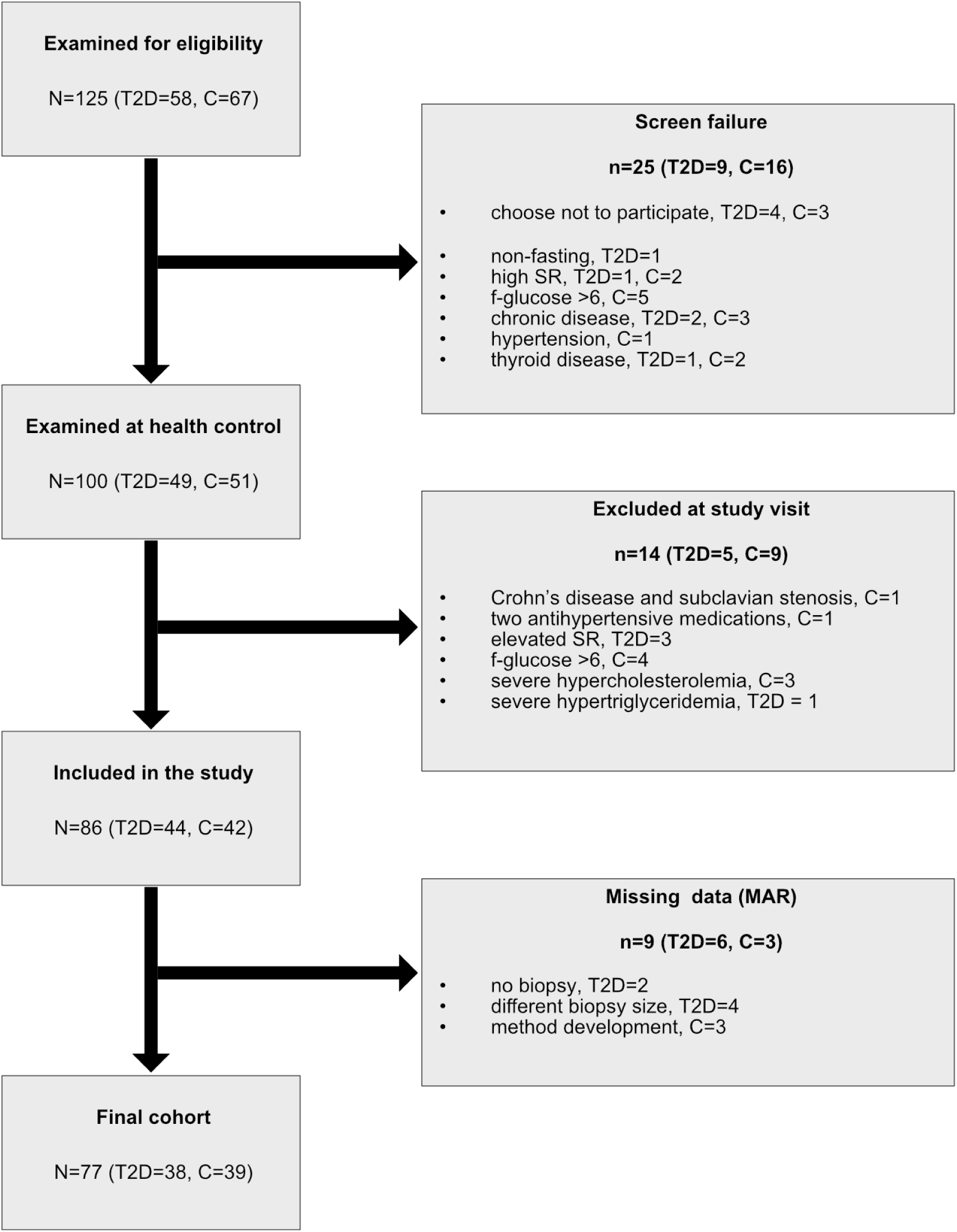
Flow diagram of participants at each stage of the study.

### CoLaus validation patient cohort

We included a nested group of 699 individuals from the CoLaus cohort for validation that has been described previously [11]. In brief, a non-stratified random sample of 35% of the Lausanne inhabitants aged 35-75 years (n=56,694) was drawn. Individuals were recruited between June 2003 and May 2006, including 6,733 participants with a participation rate of 41%. The study was conducted according to the principles expressed in the Declaration of Helsinki and approved by the Institutional Ethics Committee of the University of Lausanne. Written informed consent was obtained from all subjects.

### Preparation of tissue homogenates

Tissue homogenates were prepared using Precellys homogenizer (Bertin-technologies). Shortly, tissues were weighted on ice and transferred to the Precellys 2mL tubes with 6 beads per tube. MeOH was added to correct the concentration of the homogenates. Then 9 rounds of homogenisation at 5500 rpm, 25 seconds each round was performed at 4°C. Tissue homogenates were used for further analysis. For the skin biopsy samples, lipid extraction was performed directly in the Precellys homogenisation tubes, as described below.

### Lipidomics extraction and analysis

Lipidomics analysis was performed as described previously^43^. Briefly, extraction was performed by mixing (Thermomixer (Eppendorf), 60 minutes, 37°C) the cell pellets, plasma (100µL), or tissue homogenate with extraction buffer consisting of a mixture: methanol: methyl *tert-*butyl ether: chloroform 4:3:3 (v/v/v) and internal standards. After centrifugation (16 100 rpm, 10 minutes, 37°C), the single-phase supernatant was collected, dried under N2, and stored at -20°C. Before analysis, lipids were dissolved in 100µL of MeOH Thermomixer (Eppendorf), 60 minutes, 37°C) and separated on a C30 Accucore LC column (Thermo Fisher Scientific, 150mm x 2.1 mm x 2.6 µm) or C18 ACQUITY UPLC CSH (Waters, 150mm x 2.1 mm x 1.7µm) using gradient elution with A) Acetonitrile: Water (6:4) with 10mM ammonium acetate and 0.1% formic acid and B) Isopropanol: Acetonitrile (9:1) with 10mM ammonium acetate and 0.1% formic acid at a flow rate of 260µL/minute using Transcend UHPLC pump (Thermo Fisher Scientific) at 50°C. Following gradient was used: 0min 30% (B), 0.5min 30% (B), 2min 43% (B), 3.3min 55% (B), 12min 75% (B), 18min 100% (B), 25min 100% (B), 25.5min 30% (B), 29.5min 30% (B). Eluted lipids were analysed by a Q-Exactive plus HRMS (Thermo Fisher Scientific) in positive and negative modes using heated electrospray ionisation (HESI, Shealth gas flow rate=40, Aux gas flow rate=10, Sweep gas flow rate=0, Spray voltage=3.50, Capillary temperature=320°C, S-lens RF level=50.0, Aux gas heater temperature=325°C). MS2 fragmentation spectra were recorded in data-dependent acquisition mode with top10 approach and constant collision energy 25eV. 140 000 resolution was used for full MS1 and 17 500 for MS2. Peak integration was performed with TraceFinder 4.1, Skyline daily and Compound Discoverer 3.3 (Thermo Fisher Scientific). Lipids were identified by predicted mass (resolution 5ppm), retention time (RT) and specific fragmentation patterns using in house made, Lipidcreator and MSDIAL compound databases. Next, lipid concentrations were normalised to the internal standards (one per class), wet weight, volume and for the cell measurements to the total protein amount.

List of internal standards:

- D5-1-desoxymethylsphinganine (m17:0, 860476, Avanti Polar Lipids) 100pmol/sample
- D7-Sphinganine (d18:0, 860658, Avanti Polar Lipids) 100pmol/sample
- D7-Sphingosine (d18:1, 860657, Avanti Polar Lipids) 100pmol/sample
- D7-Sphingosine-1-phosphate (d18:1, 860659, Avanti Polar Lipids) 50pmol/sample
- 1-deoxydihydroceramide (m18:0/12:0, 860460P, Avanti Polar Lipids) 100pmol/sample
- 1-deoxyceramide (m18:1/12:0, 860455, Avanti Polar Lipids) 100pmol/sample
- Dihydroceramide (d18:0/12:0, 860635, Avanti Polar Lipids) 100pmol/sample
- Ceramide (d18:1/12:0, 860512, Avanti Polar Lipids) 100pmol/sample
- SM (d18:1/12:0, 860583, Avanti Polar Lipids) 100pmol/sample
- Glucosylceramide (d18:1/8:0, 860540, Avanti Polar Lipids) 100pmol/sample
- SPLASH lipidomics standard (330707, Avanti Polar Lipids) 2.5µL/sample

Dilutions of pooled quality control samples (QC 100%, QC 75%, QC 50%, QC 25%, QC 10%) were analysed regularly during the measurement. CVs and RSQs of individual lipids species were calculated using the quality control dilution curves, and lipids not matching overall CV < 20% and RSQ < 0.9 were excluded from the dataset.

To ensure a correct identification of all 1-deoxySL cell extracts supplemented with isotopically labelled D3-1-deoxyLCB 18:0;1 (0.25µM, 24 hours, 860474, Avanti Polar Lipids) was used as an external standard. Supplemented 1-deoxySL were tracked by M+D3 isotopic enrichment and used for RT prediction.

Transitions used for the identification of Sphingolipids and 1-deoxySphingolipids:

1-deoxyCeramides and 1-deoxydihydroCeramides:

[M+H]^+^ → [M+H – H_2_O]^+^, [M+H]^+^ → [M+H – H_2_O - FA]^+^

Ceramides and dihydroCeramides:

[M+H]^+^ → [M+H - H_2_O]^+^, [M+H]^+^ → [M+H - H_2_O - FA]^+^,

[M+H]^+^ → [M+H - 2xH_2_O - FA]^+^

HexosylCeramides:

[M+H]^+^ → [M+H - Hexosyl]^+^, [M+H]^+^ → [M+H - H_2_O - FA - Hexosyl]^+^,

[M+H]^+^ → [M+H - 2xH_2_O - FA - Hexosyl]^+^

Sphingomyelins

[M+H]^+^ → [PO4-Choline]^+^, [M+H]^+^ → [M+H - H_2_O - FA – PO4-Choline]^+^,

[M+H]^+^ → [M+H - 2xH_2_O - FA - PO4-Choline]^+^

Sphingosine-1-phosphate

[M+H]^+^ → [M+H - 2xH_2_O – PO4]^+^, [M+H]^+^ → [M+H - H_2_O - PO4]^+^

Free canonical long chain bases (Sphingosine and Sphinganine):

[M+H]^+^ → [M+H - H_2_O]^+^, [M+H]^+^ → [M+H – 2xH_2_O]^+^

Free atypical long chain bases

[M+H]^+^ → [M+H - H_2_O]^+^

Note: FA represents corresponding fatty acyl.

### Total long chain base analysis

Total long chain bases analysis was performed using acid/base hydrolysis. Briefly, 100µL of plasma was extracted using MeOH (500µL) containing internal standards. The mixture was mixed at 37°C for 60 minutes (Thermomixer (Eppendorf)) followed by centrifugation at 16 100 rpm for 10 min. Afterwards, the supernatant was collected, and 75µL of HCl (32%) was added, followed by vortexing. Next, the mixture was kept at 65°C for 15 hours. Then, the mixture was neutralised using 100µL of KOH (10M) and two-phase extraction using CHCl3 and aq. NH4OH (1.5N) was performed. The CHCl3 phase was collected, dried under N2, and stored at -20°C until further analysis. Dried-free long-chain bases were redissolved in 50µL of MeOH (70% with ten mM ammonium acetate), and 10µL was used for an LC-MS/MS analysis.

List of internal standards:

D5-1-desoxymethylsphinganine (m17:0, 860476, Avanti Polar Lipids) 200pmol/sample

D7-Sphinganine (d18:0, 860658, Avanti Polar Lipids) 200pmol/sample

D7-Sphingosine (d18:1, 860657, Avanti Polar Lipids) 200pmol/sample

### Untargeted long chain base LC-MS/MS analysis

Free long chain bases were separated on a RP18 HPLC column (INTERCHIM UPTISPHERE C18-ODB, 125mm x 3.0mm x 3µm) using gradient elution with A.) 50% MeOH with 10mM ammonium formate and 0.2% formic acid and B.) MeOH at a flow rate of 600µL/minute using Transcend UHPLC pump (Thermo Fisher Scientific) at 50°C. The elution was carried under following gradient: 0min 35% (B), 9min 65% (B), 9.5min 100% (B), 11.50 min 100% (B), 12min 35% (B), 13min 35% (B). Eluted free long chain bases were ionised by heated electron spray ionisation (Shealth gas flow rate=55, Aux gas flow rate=15, Sweep gas flow rate=3, Spray voltage=3.50, Capillary temperature=275°C, S-lens RF level=68.0, Aux gas heater temperature=450°C) and analyzed by Q-Exactive plus HRMS (Thermo Fisher Scientific) in positive mode. MS2 fragmentation spectra were recorded in data-dependent acquisition mode with top10 approach and constant collision energy 25eV. 140 000 resolution was used for full MS1 and 17 500 for MS2. Peak integration was performed with TraceFinder 4.1 and Skyline. Data were normalised to ISTD and total protein amount and isotopically corrected.

### Targeted long chain base LC-MS/MS analysis

Free long chain bases were separated on a RP18 HPLC column (INTERCHIM UPTISPHERE C18-ODB, 125mm x 3.0mm x 3µm) using gradient elution with A.) 50% MeOH with 10mM ammonium formate and 0.2% formic acid and B.) MeOH at a flow rate of 600µL/minute. Eluted free long chain bases were ionised by electrospray ionisation in positive mode and analysed in multiple reaction monitoring (MRM) mode using QTRAP 6500+ (SCIEX) mass spectrometer. Peak integration was performed using Skyline daily software.

### Amino acid extraction and analysis

Amino acids were extracted from 10µL of plasma or tissue homogenate by mixing with 180µL of ice-cold methanol containing 1nmol of stable isotope-labelled amino acids (0.5nmol for Cystine) (MSK-A2-1.2, Cambridge isotope laboratories). Samples were incubated at -20°C for 30 minutes, followed by centrifugation at four °C (16 100*g,* 10 minutes). The supernatant was collected and dried under a stream of N2 and stored at -20°C until further analysis. Dried extracts were redissolved in 100µL of 0.1% acetic acid, and 5µL was used for an LC-MS/MS analysis. Briefly, amino acids were separated on a reverse-phase C18 column (EC 250/2 NUCLEOSIL 100-3 C18HD, length 250 mm, internal diameter 2 mm, Macherey-Nagel) using gradient elution with A.) 0.1% Formic acid in water B.) 100% acetonitrile. The flow rate was held constant at 0.2 mL/min at 40°C. Eluted amino acids were ionised by electrospray ionisation in positive mode and analysed in multiple reaction monitoring (MRM) modes using QTRAP 6500+ (SCIEX) mass spectrometer. Peak integration was performed using Skyline daily software.

### Proteomics analysis

#### Trypsin digestion in denaturing conditions

The protein extracts from individual skin biopsies were prepared from the dried protein residuum after lipidomics extraction by mixing with Urea (8M) in TRIS (100mM) (Thermomixer (Eppendorf), 10 minutes, 70°C). After centrifugation (16 100 rpm, 15 min) the supernatant was collected and protein content was determined using Bradford assay. Then the protein aliquotes were transferred in equivalent volumes containing 100 µg of proteins to a 96-well plate. Reduction of proteins was performed using 5 mM tris(2-carboxyethyl)phosphine hydrochloride at 37 °C for 45 minutes. Alkylation was carried out using 40 mM iodoacetamide, followed by incubation at room temperature in the dark for 30 minutes. Subsequently, samples were diluted with four volumes of 100 mM ammonium bicarbonate and digested with lysyl endopeptidase and trypsin at an enzyme-to-substrate ratio of 1:100, at 37 °C for 16 hours. Digests were acidified by adding 50% formic acid (FA) to a final concentration of 2%. Peptides were desalted using a 96-well C18 MACROspin plate with a capacity of 10-100 µg, following the manufacturer’s instructions. After drying in a centrifugal vacuum concentrator, samples were resuspended in loading buffer containing 3% acetonitrile (ACN) and 0.1% FA. The iRT kit (Biognosys AG) was added to all samples as instructed by the manufacturer.

#### Mass spectrometry data acquisition

Peptide digests were analyzed in data-dependent acquisition (DDA) and data-independent acquisition (DIA) modes on a Thermo Scientific Orbitrap Eclipse Tribrid mass spectrometer (Thermo Fisher) equipped with a nanoelectrospray ion source and coupled to an EASY-nLC 1200 system (Thermo Fisher Scientific). Peptides were loaded onto a 50 cm × 0.75 μm i.d. analytical column packed in-house with 1.9 μm C18 beads (Dr. Maisch Reprosil-Pur 120) and separated by a 120 min linear gradient at a flow rate of 300 nL/min with increasing buffer B (95% acetonitrile in 0.1% formic acid) from 3% to 30%. For DDA, a full MS1 scan was acquired over a mass range of 350–1400 m/z at a resolution of 120,000 with an AGC target of 200% and an injection time of 100 ms. DDA - MS2 spectra were acquired at a resolution of 30,000 with an AGC target of 200% and an injection time of 54 ms. To maximize parallelization, a duty cycle time was 3 s. For DIA, a full MS1 scan was acquired between 350 and 1100 m/z at a resolution of 120,000 with an AGC target of 200% and an injection time of 100 ms. Forty-one variable-width windows (Supplementary Table 1) were used to measure fragmented precursor ions. DIA–MS2 spectra were acquired at a resolution of 30,000 with an AGC target of 400% and an injection time of 54 ms. The normalized collision energy was set to 30.

#### Mass spectrometry data analysis

Prior to DIA spectra processing, a spectral library was generated using the library generation functionality of Spectronaut 15 (version 15.5.211111.50606, Biognosys AG) using the default settings with minor adaptations ^44^. In brief, the DIA and DDA files were searched against the human UniProt FASTA database (updated 2020-03-20), the MaxQuant contaminants FASTA database (245 entries), and the Biognosys’ iRT peptides FASTA database. Digestion enzyme specificity was set to Trypsin/P with specific cleavage rules. The minimum allowed peptide length was set to 5 amino acids with a maximum of two missed cleavages per peptide. Carbamidomethylation of cysteine was considered a fixed modification, and acetylation (protein N-terminus) and oxidation of methionine as variable modifications. DIA spectra were further processed with Spectronaut using the default settings with a few modifications. In short, dynamic retention time extraction was applied with a correction factor of 1. The identification of peptides and proteins was controlled by the false discovery rate (FDR) of 1%. The machine learning algorithm and Q-value calculations were run across the entire experiment. Protein quantification was carried out using a label-free quantification approach on mean peptide quantities. Protein quantification included only proteotypic peptides and global median normalization was applied.

#### Proteomic analysis of skin biopsies

The protein report generated in Spectronaut was processed using an in-house R script in R Statistical Software (version 4.2.0; R Core Team 2021) with the *protti* package ^45^. In brief, median-normalized protein abundances were log_2_-transformed and further used for statistical testing of differentially abundant proteins between healthy control and disease samples using an empirical Bayes moderated t-test as implemented in the *limma* package (version 3.52.4). The resulting p-values were adjusted by multiple testing using the Benjamin-Hochberg method. The output of the statistical analysis was filtered using the following cutoffs: q-value < 0.05 and |log2 FC| > 0.75. These proteins were considered up- or down-regulated. Functional enrichment analysis of up- and down-regulated proteins, based on gene ontology (GO) terms molecular function (MF), biological process (BF), and cellular component (CC), was performed using g:Profiler with the FDR multiple testing correction method applying a significant threshold of 0.05 (q-value < 0.05). We further applied Gene set enrichement analysis (GSEA) on logFC of protein abundance, using WEBGESTALT (WEB-based Gene SeT AnaLysis Toolkit, webgestalt.org), parsed by KEGG database. Pathways were plotted according to their enrichment score. STRING analysis of upregulated proteins (logFC>0.75, p.adj.value<0.05) was performed using String-db.org with a highest confidence score of interaction= 0.9 and subsequently MCL clustering was performed (inflation parameter = 3).

#### Definition of variable windows for DIA–MS measurements

**Definition of variable windows for DIA–MS measurements.**
| Window | Mass-to-charge ratio ($m/z$) | Charge ($z$) | Isolation window ($m/z$) |
| --- | --- | --- | --- |
| 1 | 358 | 2 | 16 |
| 2 | 373 | 2 | 16 |
| 3 | 388 | 2 | 16 |
| 4 | 403 | 2 | 16 |
| 5 | 418 | 2 | 16 |
| 6 | 433 | 2 | 16 |
| 7 | 448 | 2 | 16 |
| 8 | 463 | 2 | 16 |
| 9 | 478 | 2 | 16 |
| 10 | 493 | 2 | 16 |
| 11 | 508 | 2 | 16 |
| 12 | 523 | 2 | 16 |
| 13 | 538 | 2 | 16 |
| 14 | 553 | 2 | 16 |
| 15 | 568 | 2 | 16 |
| 16 | 583 | 2 | 16 |
| 17 | 598 | 2 | 16 |
| 18 | 613 | 2 | 16 |
| 19 | 628 | 2 | 16 |
| 20 | 643 | 2 | 16 |
| 21 | 659 | 2 | 18 |
| 22 | 676 | 2 | 18 |
| 23 | 693 | 2 | 18 |
| 24 | 710 | 2 | 18 |
| 25 | 727 | 2 | 18 |
| 26 | 744 | 2 | 18 |
| 27 | 761 | 2 | 18 |
| 28 | 778 | 2 | 18 |
| 29 | 795 | 2 | 18 |
| 30 | 813 | 2 | 20 |
| 31 | 832 | 2 | 20 |
| 32 | 851 | 2 | 20 |
| 33 | 870 | 2 | 20 |
| 34 | 889 | 2 | 20 |
| 35 | 908 | 2 | 20 |
| 36 | 929.5 | 2 | 25 |
| 37 | 953.5 | 2 | 25 |
| 38 | 977.5 | 2 | 25 |
| 39 | 1006.5 | 2 | 35 |
| 40 | 1048 | 2 | 50 |
| 41 | 1111 | 2 | 78 |

#### Cell culture

Primary skin fibroblasts, HK2 and HEK293 cells were grown in high-glucose Dulbecco’s Modified Eagle Medium (DMEM, Thermo Fisher Scientific) supplemented with 10% fetal bovine serum (FBS) and 1% Peniciline/Streptomycin (P/S) in a 5% CO2 incubator at 37°C. Human hepatocellular carcinoma cells (Huh-7) cells were grown in RPMI 1640 (Thermo Fisher Scientific) media were grown in high-glucose Dulbecco’s Modified Eagle Medium (DMEM, Thermo Fisher Scientific) supplemented with 10% fetal bovine serum (FBS) and 1% Peniciline/Streptomycin (P/S) in a 5% CO2 incubator at 37°C. Cells were tested for the Mycoplasma contamination.

#### C2C12 Myotubes differentiation

Undifferentiated C2C12 myoblasts were grown in DMEM high glucose supplemented with 20% FBS and 1% P/S in a 5% CO2 incubator at 37°C. Differentiation was initiated at 100% confluency by serum starvation using DMEM high glucose supplemented with 2% Horse serum and 1% P/S in a 5% CO2 incubator at 37°C for 14 days. The media was changed every 48 hours. Visible myotubes formation was observed after 4-5 days and around day 10 spontaneous contractions could be observed. Diabetic sarcopenia was induced by Dexamethasone treatment (1mM, 24 hours, D1756, Sigma Aldrich) as described previously^46^.

#### Silencing of AGXT, SDS, ALAT, PSAT, GLS2

siRNAs targeting human AGXT (SR300133, 3 unique 27mer siRNA duplexes, Origene), SDS (SR307515, 3 unique 27mer siRNA duplexes, Origene)., ALAT (111548, AM16708, Thermo Fischer Scientific), PSAT (112317, AM51331, Thermo Fischer Scientific), GLS2 (SR309015, 3 unique 27mer siRNA duplexes, Origene), were used to silence (knockdown). All siRNAs were mixed to an individual final concentration of 10 nM in reduced-serum media (Opti-MEM, 31985062, Thermo Fisher Scientific). Transfection was performed using Lipofectamine RNAiMAX transfection Reagent (13778150, Thermo Fisher Scientific) according to the manufacturer’s recommendations. The media was replaced after 24 hours with fresh DMEM (10% FBS, HEK293 cells) or RPMI (10%FBS, for Huh7 cells), and cells were grown in total for 72 hours before the start of labeling experiments. Knockdown efficiency was determined using qRT-PCR as described below.

#### Plasmids generation

Wild type SDS and AGXT was cloned into the pcDNA3.1 Plasmid (V79020, Thermo Fisher Scientific), following manufacturer’s recommendation (Invitrogen). Generated plasmids were confirmed by Sanger sequencing.

#### Overexpression of SDS and AGXT

HEK293 cell lines were transfected with pcDNA3.1 Plasmid containing SDS, or AGXT genes, using Lipofectamine 3000 (L3000001, Thermo Fisher Scientific). pcDNA3.1 (V79020, Thermo Fisher Scientific) was used as an Empty vector control. Transfected cells were selected by culturing in DMEM media (10% FBS) with Geneticin (G-418, Thermo Fisher Scientific, 20µg/mL) for 4 weeks.

#### Amino acid stable isotope labelling assay

For the SL labelling assay, cells were plated at 200 000 cells/mL in 6 well plates. Cells were grown for 48 hours to 70% confluence in DMEM/10%FBS/1%P/S growth media. 24 hours before harvesting, the medium was replaced with L-serine free DMEM (C4331.0500, Genaxxon Bioscience) with 10% FBS / 1% P/S and supplemented with the indicated amount of D3N15-L-serine and D4-L-alanine (DNLM-6863-PK, DLM-250-PK, Cambridge Isotope Laboratories). Cells were harvested by trypsinisation, and cell pellets were washed 2 times with cold PBS. Cell pellets were then frozen and kept at -20°C until extraction. De novo formed SL were tracked by M+D3 isotopic enrichment, due to the one deuterium loss during the SPT reaction.

#### UC13Glucose and 15NGlutamine to Sphingolipids Flux method

To assess de novo sphingolipid synthesis from glucose and glutamine, we performed metabolic flux assays using uniformly labeled UC13Glucose (CLM-1396-1, Cambridge Isotope Laboratories) and 15N-(Amine)-labeled L-glutamine (15N-Glutamine, HY-N0390S, MedChemExpress). For the UC13Glucose tracing, cells were seeded in 6-well plates at a density of 200 000 cells per well and cultured for 48 hours in complete medium.. Following this period at 70% confluency, the culture medium was replaced with glucose-free, serine-free medoa DMEM (C4332,0500, Genaxxon bioscience). supplemented with 10 mM UC13Glucose and additional treatments as indicated in the respective figure panels. After the incubation time points specified in the figures, cells were harvested using 0.25% Trypsin-EDTA, washed twice with ice-cold phosphate-buffered saline (PBS), and used for further analysis..

UC13Glucose forms 3C13Alanine and 3C13Serine. During the SPT reaction C1 carbon of the amino acid is released as 13CO2. Therefore, UC13Glucose derived SL and 1-deoxySL would end up with an enrichment of 2C13 assuming they are coming from the 3C13Alanine and 3C13Serine. It should be noted that other isotopologues of these labelled amino acids are formed but would result in an enrichment of 1C13 in SL backbond. Complex SL have usually high number of carbons (up to 44 carbons) which makes isotopic enrichment of +2C13 difficult to measure due to a very low isotopic enrichment. Therefore, we developed an UC13Glucose flux method combining lipid extraction combined with acid hydrolysis to release the long chain base (18 carbons) and therefore increase relative UC13Glucose enrichment on at the long chain base.

For 15N-glutamine labeling experiments, glutamine-free DMEM was used and supplemented with 10mM 15N-glutamine or unlabeled glutamine. Experimental conditions, including co-treatment with the SPT inhibitor Myriocin or other modulators, are detailed in the figure legends. After incubation, cells were harvested and washed as described above.

#### Combined UC13Glucose BSA/D4-Sodium palmitate Sphingolipid flux method

Huh7 cells were starved with reduced FBS (1%) and treated with high concentration BSA/D4- Sodium Palmitate complexes for 16 hours prior to initiation of UC13Glucose to SL flux assay. BSA/D4- Sodium palmitate complexes were prepared as follows. First, 10µmol of D4-Palmitic acid (901412, Sigma Aldrich) was weighted into an Eppendorf tube and EtOH (100µL), H_2_O (300µL) and 0.1M NaOH (100µL) were added. Next, the mixture was heated at 75°C and mixed for 2 hours (Thermomixer (Eppendorf)). Afterwards, 500µL of H_2_O were added (yielding total 10mM concentration). BSA complexes were prepared by the addition of 100µL of fatty acid sodium salts to 900µL BSA (Essentially fatty acids free (Thermo Fisher Scientific)) in DMEM (6.66mg/mL) and shaking at 30°C for 1 hour ((Thermomixer (Eppendorf)) and used directly for isotope labelling as described above.

#### qRT-PCR

For qRT-PCR, cells were harvested using trypsinisation as described above. Cell pellets or tissue homogenates were lysed using TRIzol reagent and RNA was extracted using the phenol/chloroform method following manufacturer‘s instructions. RNA was purified using Ethanol and concentration/purity was determined using NanoDrop (Thermo Fisher Scientific). Reverse transcription was performed using Maxima Reverse Transcriptase following manufacturer‘s instructions (EP0742, Thermo Fisher Scientific).

qPCR reactions were performed using SYBR Green qPCR mastermix. Absolute concentration of mRNA was calculated from the dilution curve (1/10, 1/100, 1/1000 and 1/10000 dilution) of an adequate plasmid. GAPDH or 18S were used as an in-house gene loading control.

### Protein determination

For the protein content normalization, cell pellets remaining after the lipid extraction were dissolved in Urea (8M) containing 1% 2-Mercaptoethanol and mixed at 800rpm at 90°C (Thermomixer (Eppendorf)) for 20 minutes. Next, protein extracts were snap frozen using dry ice, and the whole process was repeated 3 times. After centrifugation (16 100 rpm, 10 minutes), supernatants were collected and used for further analysis. Total protein amounts were determined using Bradford assay (Bio-Rad, 1:5 dilution) according to the manufacturers’ recommendation. External calibration curve of Bovine serum albumine was used to calculate total amounts of protein.

### Fluorescence microscopy

For the fluorescence microscopy cells were grown in 96-well plates in the incubator at 5% CO_2_ at 37°C. Indicated treatments were added by exchange of the media. After the given time (indicated in the figures), cells were washed 3 times with PBS and fixed using 4% PFA for 30 minutes. Next, cells were washed 3 times with PBS and then incubated for 1 hour with DAPI (D9542, Sigma-Aldrich, 1 µM in PBS) and Phalloidin-665 (18846, Sigma-Aldrich, 0.5 µM in PBS). Further, cells were washed with PBS and imaged immediately. Images were acquired using a fluorescence microscope (Olympus IX81) with a motorized stage at the 20x magnification.

### Alanine aminotransferase (ALAT) activity assay

The ALAT activity was determined using the commercially available kit (ab105134, Abcam) following the manufacturer’s instructions. Briefly, 50mg of liver tissue was homogenised using Precelysis tissue homogeniser in a cold ALT lysis buffer. Next, extracts were diluted 1:2000 in ice-cold ALT lysis buffer and activity was measured using the protocol for fluorimetric determination. Two technical replicates per each liver sample were measured, and the average was used for data analysis.

### Mouse experiments

C57BL/6J mice were fed an obesity inducing diet with w/60% energy from fat (#C1090-60, Altromin Spezialfutter GmbH & Co.KG, Lage, Germany) or respective control diet for 16 weeks. At the time of analysis, mice were obese and hyperinsulinemic, and featured pathological glucose and insulin tolerance tests. Plasma and tissue samples were obtained following an overnight fast and stored at - 80°C. This procedure was formally approved by and conducted in accordance with the established institutional guidelines of the Landesamt für Natur, Umwelt und Verbraucherschutz NRW, Germany.

Male obese and hyperglycemic db/db BKS (The Jackson Laboratories/JAX #000642) and heterozygous control animals were acquired from Charles River (local distributor) and acclimatized for one week before sacrifice. Mice were maintained on a 12 h light/dark cycle (lights on from 6:00 to 18:00) and had ad libitum access to tap water and a standard rodent chow (58% carbohydrates, 33% protein, 9% fat, R/M−H Extrudat, ssniff Spezialdiäten GmbH, Soest, Germany). Plasma and tissue samples were rapidly frozen in liquid nitrogen and stored at -80°C, until shipment to Zurich surrounded by dry ice. Animal procedures were approved by the Department for Environment and Consumer Protection of North Rhine-Westphalia, Germany (LANUV; # 81-02.04.2020.A086) and the DDZ Institutional Animal Welfare Committee.

### Rat experiments

The rats were randomized into two experimental groups: a control group (n = 10) and a diabetic group (n = 15). Type 1 diabetes mellitus (T1DM) was induced in the diabetic cohort via a single intraperitoneal administration of streptozotocin (STZ, 55 mg/kg; Sigma-Aldrich, St. Louis, MO, USA) following an overnight fasting period. STZ was freshly prepared in 0.1 mol/L citrate buffer (pH 4.5) immediately prior to injection. Glycemic status was assessed 72 hours post-injection by measuring blood glucose levels from the tail vein using an Accutrend Plus glucometer (Roche, Switzerland). Animals with a fasting blood glucose concentration of 12 mmol/l were considered diabetic. After six weeks of follow-up, animals were sacrificed via a controlled asphyxiation in a carbon dioxide chamber. Hepatic tissues were immediately excised, snap-frozen in liquid nitrogen, and stored at −80°C for subsequent analyses. Blood samples were collected from the vena cava inferior into sterile tubes, allowed to clot at room temperature for 1 hour, and subsequently centrifuged (10 min, 1100–1300 × g, 4°C). The resulting sera were aliquoted and stored at −80°C until further analyses.

### STZ L-serine supplementation

In accordance with the 3R principle, we used tissues from the previously published study^22^. Briefly, male Sprague-Dawley rats (180-200 g, Charles River) were used. The animals had access to food and water supplies ad libitum. Diabetes was induced in overnight-fasted rats by a single intraperitoneal injection of 60mg/kg streptozocin (STZ) (Sigma-Aldrich) dissolved in a vehicle (sodium citrate buffer, pH 4.5). Control rats were injected with the vehicle alone. L-ser supplementation in STZ rats was performed by access to a L-ser-enriched diet (5% L-ser). At the end of the study, animals were sacrificed; tissues were dissected and immediately frozen in liquid N_2_ and stored at -80°C until further use.

### Bortezomib treatment

The effect of Bortezomib was tested in a rat model as described before ^47^. Briefly, Bortezomib (LC Laboratories, Woburn, MA) was prepared in a vehicle solution composed by absolute ethanol/tween80/saline (5%/5%/90%) and administered intravenously via the caudal vein. For chronic exposure, rats received intravenous Bortezomib at a dose of 0.20 mg/kg, administered three times per week over a period of 8 weeks. Control animals received vehicle only. Plasma samples were obtained following an overnight fast and stored at -80°C. At the end of the study, animals were sacrificed.

### Data analysis, statistics and Figures

All the data analysis and figure preparations were performed using GraphPad Prism 9.5.1 (Bar graphs, Violin plots, Scatter plots), R (Differential analysis, Correlation analysis), Metaboanalyst 5.0 (Differential analysis, Volcano plots), Stata (Correlation plots) and Excel. Statistical analysis was performed in GraphPad Prism 9.5.1. Illustrations were made using BioRender webpage (BioRender.com). Final figures were prepared using Affinity Designer 2.

## Author contributions

MA designed and performed the majority of experiment. BP, HJ and AB characterised the T2D cohort, collected plasma samples and skin biopsies; HA and MA performed the proteomics analysis; AC, OA and GL performed the rats samples and collected tissue samples; BL, BB prepared mice tissue samples; YE prepared tissues homogenates; EM performed the qPCR analysis; AC and AZ performed siRNA knockdowns, overexpressions and corresponding assays, AC performed C2C12 differentiation and corresponding analysis, RJ and KS isolated HDL and LDL lipoprotein populations; HT helped with the data analysis and co-authored the manuscript.

## Acknowledgements

This project was supported by the Swiss National Science Foundation (SNF 310030_215134) and by the SNF under the frame of the European Joint Program on Rare Diseases (EJP RD+SNF 32ER30_187505).

“This publication is based upon work from COST Action 19105-Pan-European Network in Lipidomics and EpiLipidomics (EpiLipidNET) supported by COST (European Cooperation in Science and Technology)”. We are also grateful to Peter Vollenweider for sharing the CoLaus data.

